# Structural basis for start codon selection by eIF5-mimic proteins

**DOI:** 10.64898/2025.12.28.696766

**Authors:** Ximena Zottig, Chun-Ying Huang, Zahra Seraj, Nikolaus Grigorieff, Andrei A. Korostelev

## Abstract

Start codon selection extensively regulates the cellular proteome and is implicated in cancer and other diseases. The eIF5-mimic proteins (5MPs: 5MP1/*BZW2* and 5MP2/*BZW1*) restrict non-AUG start codon usage and thereby reprogram proteoform expression from mRNAs with alternative start sites, such as the oncogenic c-Myc. Here, using *in extracto* cryo-electron microscopy (cryo-EM) and biochemical assays, we report how 5MP represses translation dependent on the sequence context at start codons. Cryo-EM structures of 48S pre-initiation complexes (PICs) reveal 5MP1 bound at the A site of the small ribosomal subunit. 5MPs act as initiation factors that stabilize an open PIC conformation, promoting mRNA scanning and inhibiting initiation at suboptimal start codons. We propose to designate 5MP as eIF1S (eukaryotic Initiation Factor 1 Scanning). These findings reveal the structural basis for proteome modulation by 5MPs and may help correct dysregulated translation in disease.

## Main

Translation initiation is a major regulatory step of gene expression, and its dysregulation contributes to a wide range of human diseases, including cancer (*1–3*). During initiation, the small ribosomal 40S subunit, together with initiation factor proteins (eIFs) and initiator tRNA^Met^, collectively termed the 48S pre-initiation complex (PIC), scans the 5ʹ untranslated region (UTR) of an mRNA until it encounters the initiation (start) codon (*4–7*). At the start codon, the small subunit associates with the large 60S subunit to commence protein synthesis. Although the AUG triplet is the predominant start codon, recent ribosome profiling and other studies revealed extensive initiation from non-AUG codons, notably from upstream open reading frames (ORFs), expanding proteome complexity (*8, 9*). A recent study has shown that thousands of ORFs lie outside annotated coding sequences and encode for microproteins, strengthening the case for studying upstream ORFs and mechanisms controlling their translation (*10, 11*). Moreover, switching between AUG and non-AUG codons within a single mRNA can control the output of distinct proteoforms (*12–14*). For example, the increased initiation stringency, favouring AUG codons, in *c-Myc* and other mRNAs enhances translation of tumor-promoting transcripts (*15–19*). Likewise, AUG codons in suboptimal contexts (*i.e.*, sequences near the start codon, also known as Kozak sequences, as detailed below) are regulated by initiation stringency, further shaping the proteome (*20*).

The eIF5-mimic proteins 5MP1 and 5MP2 are similar and conserved across eukaryotes, except in yeasts and nematodes (*21*), are encoded by the *BZW2 and BZW1* genes in humans, respectively, and regulate the selection of AUG start codons by suppressing initiation at non-AUG codons (*22–24*). Their C-terminal W2/HEAT3 domain is homologous to that of eIF5, an initiator factor promoting 48S progression toward subunit joining upon start-codon recognition, but the N-terminal domain bears no homology with eIF5. 5MP is thought to antagonize eIF5 and bind the W2-interacting eIF2β subunit of eIF2, whose rearrangement is critical for 48S PIC switching from the scanning to start-codon recognition mode (*22–26*). 5MP1 and 5MP2 sequences and cellular phenotypes are similar, indicating that they perform similar molecular roles while differing in cell/tissue expression, localization, and/or other aspects of cellular control (*21–23*). 5MP2 mRNA and protein are highly expressed in most tissues, achieving levels comparable to or exceeding those of core initiation factors (*e.g.,* eIF1A and eIF5), whereas 5MP1 is predominantly enriched in heart and muscle tissues (GTEx; Human Protein Atlas). Elevated expression of 5MP correlates with tumorigenesis, metastasis, and poor prognosis; furthermore, 5MP1 is thought to contribute to the malignant phenotype in colorectal cancer (*19, 27–29*). The oncogenic activity of 5MPs is attributed, at least in part, to the direct upregulation of the transcription factor c-Myc, a well-characterized oncogene that can drive malignant progression (*30*). Specifically, increased 5MP levels suppress translation of the long c-Myc isoform from a CUG start codon and upregulate expression of a shorter, more tumorigenic isoform from a downstream AUG codon (*19*). Furthermore, the critical roles of 5MP in start-site selection have been documented in embryonic development, neuronal homeostasis, and neurodegeneration (*23, 31, 32*).

Despite its critical cellular functions and therapeutic relevance, the molecular basis of 5MP action remains unknown due to a lack of structural data. Using *in extracto* cryo-EM—a method we recently developed for high-resolution visualization of cell lysates—we identified 5MP among endogenous translation initiation factors bound to the 40S subunit (*33*). In the resulting 2.7-Å structure of a 48S pre-initiation complex, 5MP resides in the A site and stabilizes the open, scanning-competent conformation of the PIC through interactions with the 40S subunit, initiator tRNA^Met^, and the initiation factors eIF2β and eIF1A. Our mutational analyses reveal that the N-terminal tail in the mRNA tunnel plays a critical role in the stringency of codon selection. Comparison of numerous start-site contexts shows that 5MPs act by preventing commitment to suboptimal initiation sites, resulting in AUG-initiated isoforms of *c-Myc* and other mRNAs. Together, these findings establish 5MP as a critical initiation factor that defines start-codon fidelity and reveal the structural basis of 5MP-mediated control of translation initiation.

### 5MP binds 48S and inhibits initiation on mRNAs with non-AUG and poor Kozak-context starts

To visualize native interactions of the translation initiation machinery with endogenous cellular components, we performed cryo-EM of mammalian cell extracts combined with 2D template matching (2DTM), an approach we refer to as *in extracto* cryo-EM (*33*). 2DTM is a computational approach that we developed to identify individual particles in noisy, low-contrast environments (*34, 35*), such as cellular lysates. Traditional single-particle cryo-EM requires purifying the macromolecule(s) of interest from cellular components and/or reconstitution of a complex from a defined subset of purified factors. To allow for identification of novel cellular complexes, *in extracto* cryo-EM uses a known fragment of a molecular template (e.g., a part of a ribosomal subunit) to detect complexes of interest in the context of cellular components. This approach was recently used to resolve near-atomic-resolution structures of endogenous 80S complexes in cell lysates, demonstrating that *in extracto* cryo-EM can recover a new set of bound proteins that were inaccessible in purified or reconstituted systems (*33*). In the present work, we used the 40S body as the 2DTM template because it is present in all PICs and undergoes fewer conformational changes than other regions of the 40S subunit, such as the head. 3D classification of particles identified by 2DTM in cellular lysates revealed a class whose A site exhibited density that could not be assigned to known translation factors (Fig. 1A; fig. S1; Methods). This class corresponds to a 48S initiation complex, as indicated by the presence of initiation factors, initiator tRNA^Met^, and mRNA (fig. S1B; Methods). Backbone and side-chain tracing with ModelAngelo (*36*) in our 3 Å resolution map, followed by comparison with the AlphaFold-predicted structure library (*37*), indicated that the density can be accounted for by a 5MP protein (Fig. 1A; fig. S1D; Methods). This finding suggests that 5MPs may directly and specifically control translation initiation from AUG or non-AUG codons by binding to the A site. To corroborate this idea and further elucidate the mechanisms by which 5MP regulates start-codon selection, we purified recombinant human 5MPs and structure-guided mutants (fig. S2) for biochemical and additional cryo-EM analyses.

**Fig. 1.**
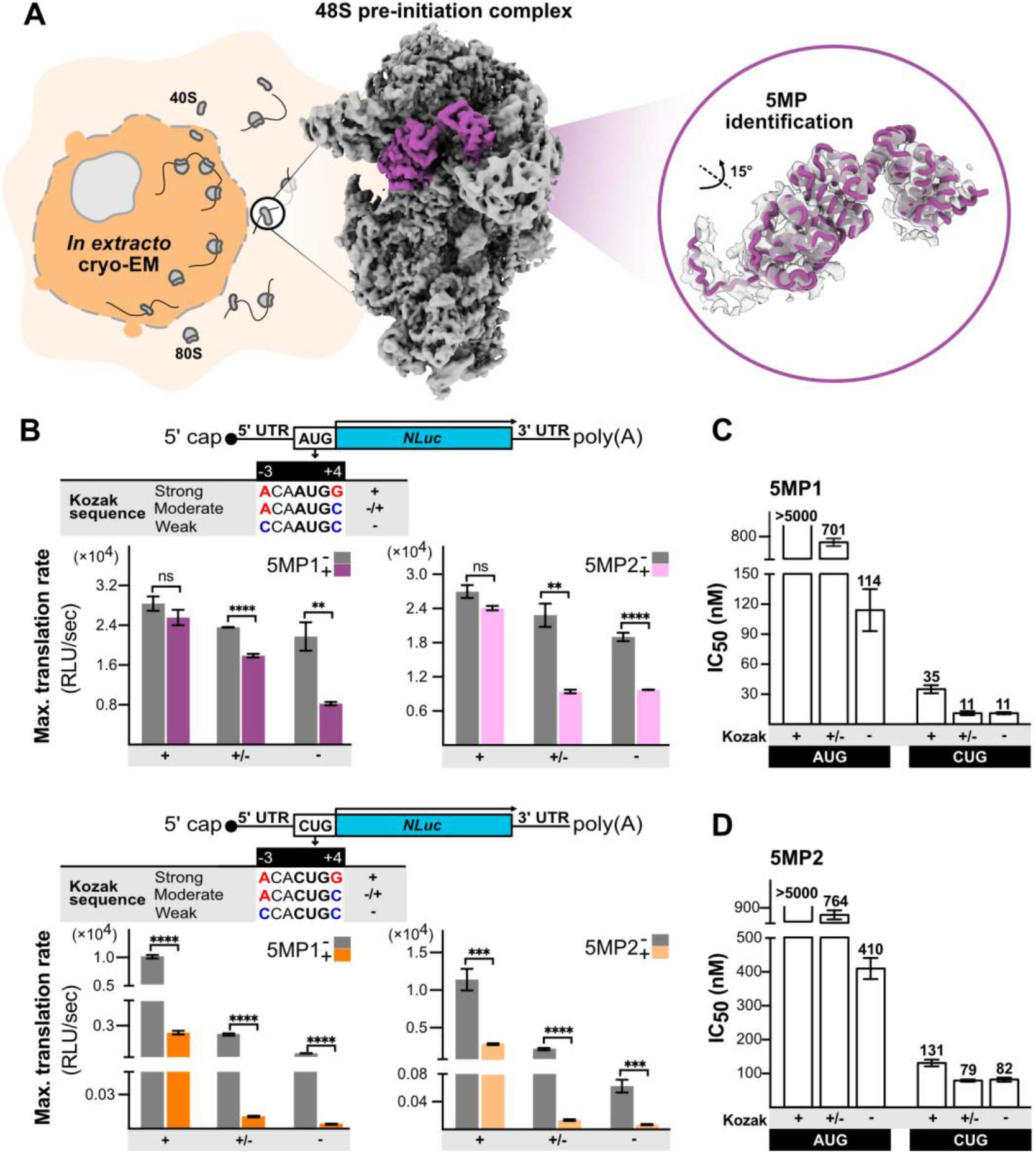
5MP proteins bind the cellular 48S pre-initiation complex and inhibit translation initiation in a Kozak sequence-dependent manner. **(A)** Schematic of *in extracto* cryo-EM and reconstruction of the 48S pre-initiation complex, highlighting unassigned density overlaid with a refined AlphaFold2-predicted model of eIF5-mimic protein (5MP) (close-up). **(B)** Schematic of mRNA constructs and effects of 5MPs at 1 μM on translation from AUG (purple) and CUG (orange) start codons in strong (+, −3A/+4G), moderate (+/−, −3A/+4C), and weak (−, −3C/+4C) Kozak contexts (mean with SEM, n=3). The statistical difference between the translation rates is shown, where ****P < 0.0001, ***P < 0.0002, **P < 0.0021, *P < 0.00332, and ns (non-significant) according to the unpaired two-tailed t-test. Left: 5MP1; Right: 5MP2. **(C)** Inhibitory concentrations (IC50) of 5MP1 (top) and **(D)** 5MP2 (bottom) for AUG and CUG initiation in strong (+, −3A/+4G), moderate (+/−, −3A/+4C), and weak (−, −3C/+4C) Kozak contexts.

To quantify 5MP-mediated regulation, we measured translation of a nanoluciferase (NLuc) reporter mRNA with an AUG or CUG start codon within strong, moderate, or weak Kozak contexts (*38, 39*). For example, a strong Kozak context with −3 purine and +4 guanosine (where +1 is the first position of the start codon) results typically in efficient translation, whereas −3 and +4 pyrimidines render the start codon inefficient (*38*). We added mRNA reporters to a rabbit reticulocyte lysate (RRL) translation system in the presence or absence of recombinant 5MPs and measured luminescence (Fig. 1B). We estimated that the endogeneous 5MP concentration in RRL is below 25 nM (fig. S2c). Translation from an AUG codon in a strong Kozak context was unaffected in the presence of 1 µM 5MPs, underscoring the robustness of canonical initiation (IC_50_ > 5 µM). By contrast, translation from moderate and weak AUG contexts was noticeably reduced in the presence of 1 µM 5MP (Fig. 1B). 5MP1 inhibited translation from a weak AUG context more efficiently than did 5MP2 (IC_50_ values of 114 nM and 410 nM, respectively) (Fig. 1C, D). As 5MP1 and 5MP2 share ∼70% overall sequence identity (Expasy; SIM alignment), including fully conserved residues mediating PIC engagement, the difference in potency may be due to different interactions with other cellular components via less conserved regions or other factors to be established (see Discussion). Translation from CUG codons—especially in moderate and weak contexts—was even more efficiently inhibited by 5MPs at low nanomolar IC_50_ values (5MP1: 35-11 nM; 5MP2: 131-79 nM; Fig. 1C, D; fig. S3A, B). Overall, the inhibitory effect of 5MP can therefore be >100-fold stronger on CUG codons than on the strong AUG codon. These findings align with previously reported results in a cellular context, indicating that the observed regulatory effect is not specific to RRL (*25*). For example, cellular levels in HEK293T cells are estimated at ∼750 nM and ∼820 nM for 5MP1 and 5MP2, respectively (*40*), well above the IC_50_ values measured for weak AUG and CUG contexts, suggesting that this inhibitory activity is physiologically relevant *in vivo*.

5MP proteins autoregulate their translation from an AUG in a conserved weak Kozak context (*24*). We therefore measured the ability of 5MP to inhibit translation from the UUUAUGA context of 5MP (*BZW2*) mRNA (fig. S3C). Indeed, 5MP1 inhibited translation from its own AUG with an IC_50_ of ∼253 nM, underscoring the ability of 5MP to repress a moderate to weak AUG context (fig. S3D, E). In cells, this feedback mechanism likely regulates 5MP concentrations, enabling differential translation from AUG and non-AUG codons across the transcriptome (*24*).

### 5MP1 regulates proteoform output by start-site selection

5MP1 drives oncogenesis, at least in part, by reprogramming expression of c-Myc from a p67 proteoform initiated at a CUG start codon to an oncogenic p64 proteoform initiated at an in-frame downstream AUG codon (*19*). The “p67” CUG start codon is in a strong Kozak context, and the “p64” AUG start codon is in a moderate Kozak context (Fig. 2A). To dissect the effects of 5MP1 on each start codon, we generated a reporter mRNA comprising the c-Myc 5′UTR and first 48 nucleotides of the p64 coding sequence fused to 3×FLAG and NLuc, and measured luminescence (Fig. 2A, B; fig. S4A) and proteoform levels (Fig. 2C, D) in the absence or presence of increasing concentrations of 5MP1. Consistent with studies in cell lines (*19, 23*), 5MP1 increased the total expression of the c-Myc construct (Fig. 2B-D) and shifted expression toward the shorter AUG-dependent proteoform by inhibiting the expression from the p67 CUG start codon (Fig. 2C, D). Mutations that disrupt the AUG or CUG codon individually confirm this trend (Fig. 2B-D; fig. S4A, B). In the absence of the AUG codon, 5MP1 suppresses CUG translation at a 1-micromolar concentration (Fig. 2C, D). By contrast, AUG initiation from its native position in the c-Myc 5′UTR is robust even at 15-micromolar 5MP1. Together, our data demonstrate that 5MP1 efficiently changes the ratio of CUG and AUG proteoforms of c-Myc at 5MP1 concentrations that are above its autoregulatory IC_50_ (fig. S3E).

**Fig. 2.**
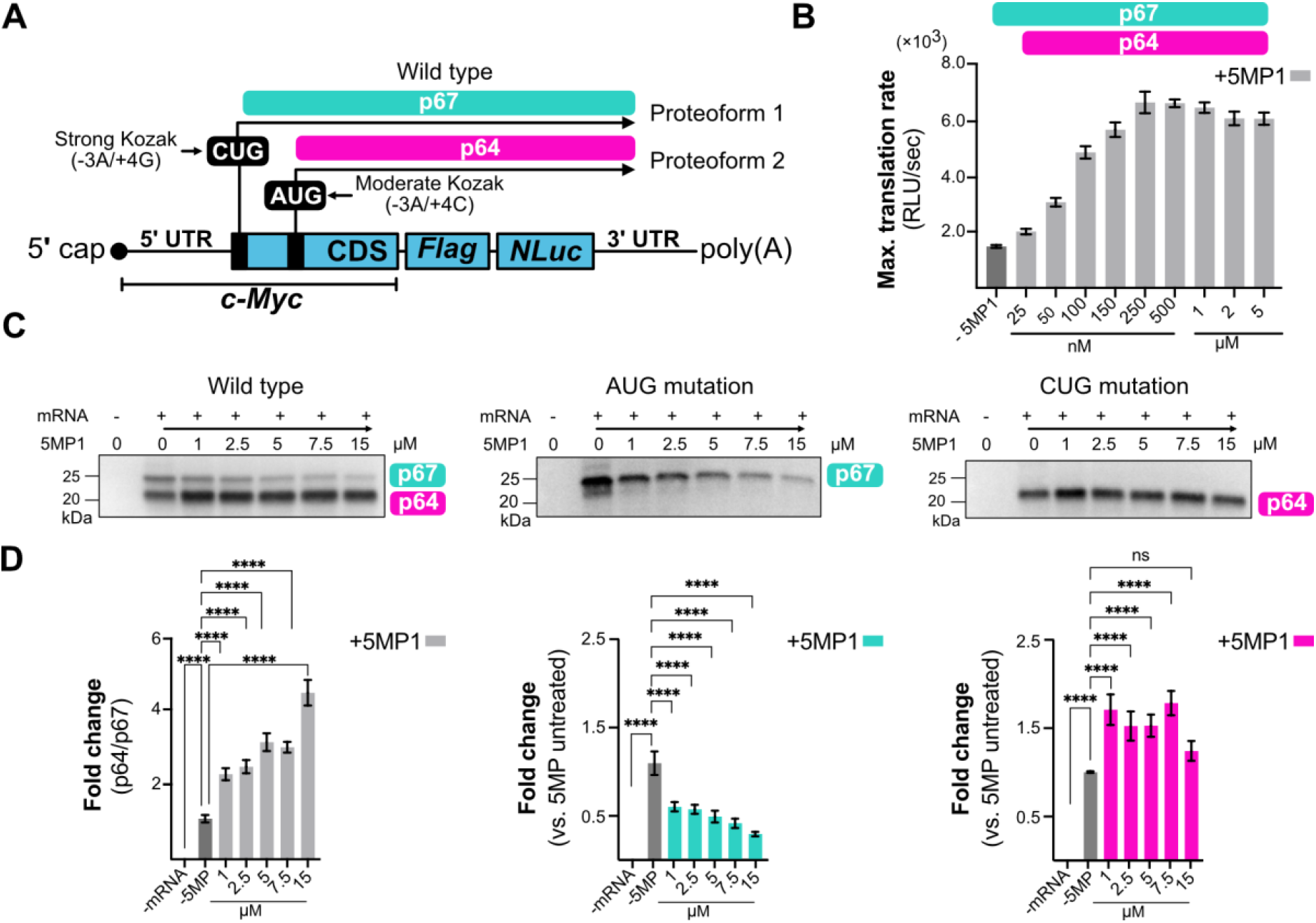
5MP1 reprograms initiation from the *c-Myc* 5ʹ UTR. **(A)** Schematic of the mRNA reporter with c-Myc 5ʹ UTR containing CUG and AUG start codons in distinct Kozak contexts. AUG-initiated p64 products are indicated in pink and CUG-initiated p67 products in green. **(B)** Maximum translation rates plotted at different 5MP1 concentrations for the c-Myc wild type reporter (see also fig. S4). Data represent mean ± SD values (n = 3). **(C)** Translation products from wild-type, AUG-mutated, and CUG-mutated constructs. Representative western blotting and **(D)** quantifications are shown: (left) fold change in proteoform ratio (p64/p67); (middle and right) p67 and p64 fold changes relative to 5MP1 untreated. Statistical significance was assessed by one-way ANOVA; ****P < 0.0001, ***P < 0.0002, **P < 0.0021, ns = not significant.

On the c-Myc untranslated region with both start codons, 5MP1 not only repressed the production of the CUG isoform but also increased the AUG isoform output (Fig. 2C, D). We wondered whether the suppression of the AUG isoform in the absence of 5MP1 might be caused by inefficient initiation on the CUG start codon, inefficient elongation between the CUG and AUG codons (*e.g.*, due to suboptimal, or “rare”, codons), or both. To distinguish between these possibilities, we generated a series of c-Myc reporter mutants and measured their translation (Fig. S4C, D). Mutating the upstream CUG while leaving the downstream AUG (Mutant 1) had no effect on translation. By contrast, mutating the AUG while retaining the upstream CUG and/or another potential start codon GUG (Mutants 2-4) reduced translation 2-fold relative to the wild-type c-Myc transcript. Converting the upstream CUG to an AUG (Mutant 5) restored translation to nearly wild-type levels, indicating that the baseline suppression of downstream AUG is primarily due to inefficient initiation at the upstream CUG and GUG codons. Furthermore, replacing the suboptimal elongation codons between the CUG and AUG start codons with synonymous optimal codons (Mutant 4) did not increase translation from the CUG codon, arguing against inefficient elongation. These findings are consistent with biochemical studies showing that AUG initiation is at least twice as efficient as initiation at near-cognate UUG and CUG start codons in the same sequence context (*41–46*). Accordingly, the less efficient initiation from the CUG-p67 start codon results in lower overall c-Myc production. Notably, the ∼400-nt long c-Myc 5’ UTR harbors nearly 20 in-frame and out-of-frame near-cognate start codons that, in the absence of 5MP1, may compete with the AUG codon for ribosome engagement (Table S1). More generally, the probability of encountering an NUG triplet at any 5’ UTR position is ∼1/16, making the scanning process on an average mRNA inefficient and prone to low-fidelity initiation without 5MP. Suppression of these competing initiation events by 5MP would be necessary for 48S to efficiently scan through to the AUG codon, thereby explaining the observed increase in AUG isoform output. In summary, our data demonstrate that when near-cognate and canonical start codons compete for ribosome engagement, 5MP1 proteins shift the isoform ratio toward the AUG-initiated isoform and increase its output.

### 5MP1 stabilizes scanning-competent 48S PIC

To investigate the structural basis of 5MP1-mediated control of initiation, we collected a cryo-EM dataset from RRL supplemented with purified recombinant human 5MP1, an mRNA bearing an AUG codon in a moderate Kozak context, and the non-hydrolyzable GTP analog Guanosine 5′-[β,γ-imido]triphosphate (GMPPNP) to capture initiation complexes with GTPase initiation factors eIF2 and/or eIF5B (*47–51*). To enable particle detection despite the low contrast of lysate micrographs, we employed 2DTM (*34, 35, 52*) using the 40S body as the search template (PDB: 7R4X; Table S2, Methods). The detected particles comprised a range of structurally distinct PICs and 80S complexes, indicating that 2DTM was mostly unbiased by the incomplete-subunit template. Cryo-EM data classification yielded predominantly 48S PICs (∼31%), containing eIF1, eIF1A, eIF2 and eIF3. Nearly half of PICs contained 5MP1, of which 37% are 48S complexes and 12% are 48S-like assemblies containing the N-terminal domain of eIF3c subunit near eIF1 but missing the rest of eIF3 (fig. S5B-D; fig. S7). Specifically, a previously unresolved segment of eIF3c forms an α-helix (Thr250 to Asp290), whose conserved FLKK motif 3 inserts into a hydrophobic pocket of eIF1 (fig. S7D), consistent with the recent report of eIF3c/eIF1 interfaces (*53*). These observations suggest a correlation between eIF3c stabilization in a PIC and 5MP1 binding (*23*). Other classes included free 40S subunits and 40S complexes containing initiator tRNA^Met^, eIF1, eIF1A, and eIF2 (fig. S5B).

A 2.7-Å reconstruction of the predominant 48S•5MP1 complex features 5MP1 at the A site (Fig. 3), placed identically to the endogenous 5MP1 detected in cell lysates described above (Fig. 1; fig. S1). Notably, cryo-EM data contain both open-head (*i.e.*, scanning-competent) and closed-head (*i.e*., codon-recognition) 48S PICs, but 5MP1 is only bound to 48S PIC in the open-head conformation (Fig. 4; fig. S5-7). In canonical open-head initiation complexes, the head is tilted ∼13° away from the body, relative to that in closed-head (codon-recognition) complexes (*54–56*). In 5MP1-bound complexes, the 40S head is tilted away by ∼15°, indicating that 5MP1 stabilizes a more widely open state and consequently features a larger mRNA tunnel compared to the canonical scanning conformation (Methods) (*54, 55*). The initiator tRNA^Met^ is positioned ∼7 Å away from its codon-engaged position, nearly identical to P_out_ conformations observed in scanning-competent 48S PICs (Fig. 4B) (*57, 58*). Although relatively continuous mRNA density is visible in the mRNA tunnel within the 5MP1-bound complexes (fig. S8), individual nucleotides are not resolved near the P site, consistent with the heterogeneity observed in scanning 48S complexes (Fig. 4C; fig. S8) (*4, 7*). This contrasts with a well-resolved AUG codon in our 2.8-Å reconstruction of codon-recognition closed-head 48S complexes lacking 5MP1, which is consistent with observations in previous studies of canonical AUG initiation (Fig. 4, C; fig. S8) (*4, 7*).

**Fig. 3.**
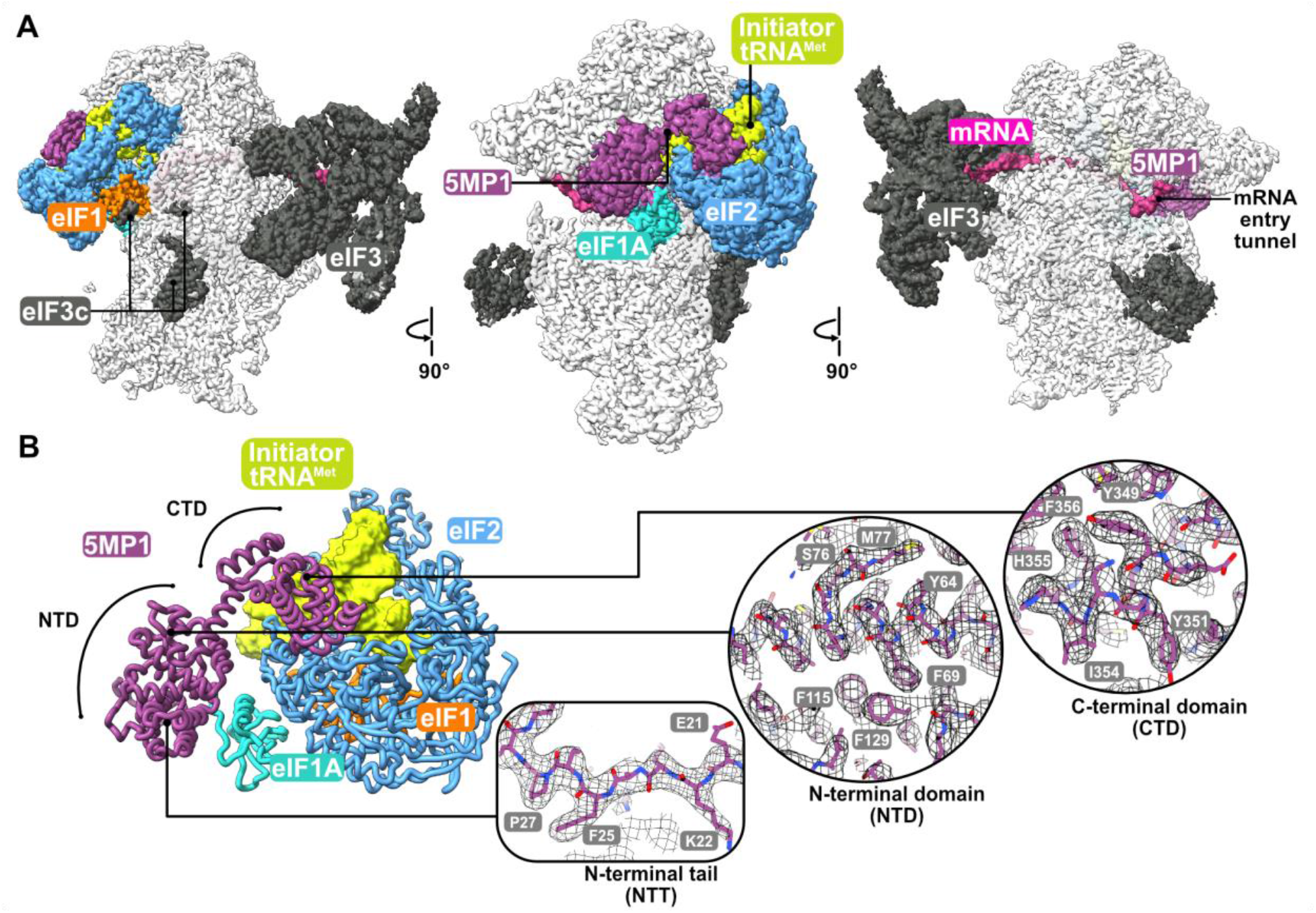
Cryo-EM of 5MP1 bound to 48S in RRL. **(A)** A 2.7-Å cryo-EM map of the 48S•5MP1 initiation complex, shown in different orientations. Segmented densities show: 5MP1 in purple, eIF2 in blue, eIF1 in orange, eIF1A in turquoise, and the initiator tRNA^Met^ in green. The middle view shows 5MP1 binding at the A site. The left and right panels show the intersubunit and solvent sides of the 48S initiation complex. **(B)** 5MP1 interactions with initiation factors and initiator tRNA^Met^. Insets show the N-terminal tail (NTT), N-terminal domain (NTD), and C-terminal domain (CTD); cryo-EM density is shown as mesh.

**Fig. 4.**
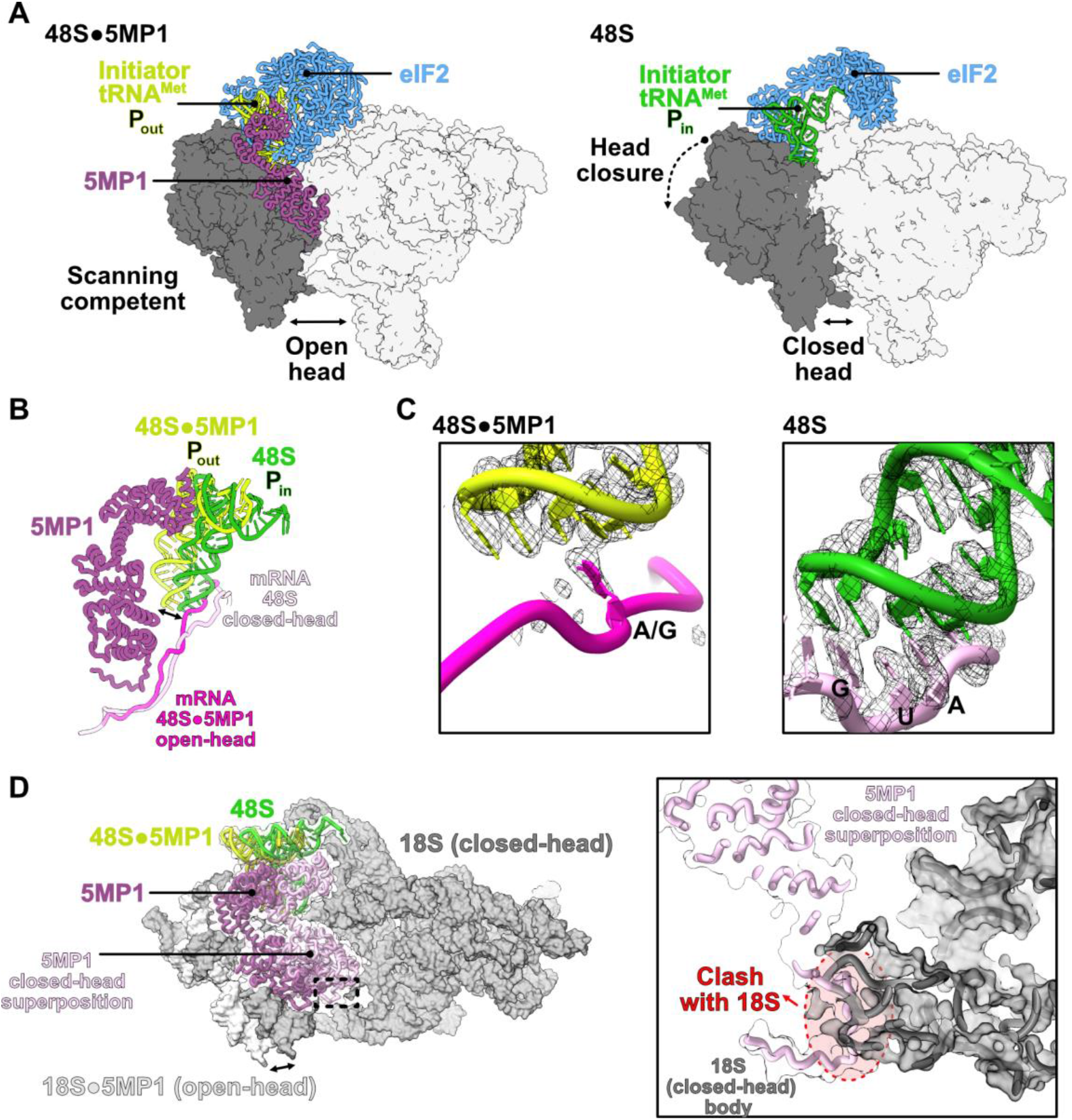
5MP1 promotes the open-head conformation of the 48S complex. **(A**) The 48S complex containing 5MP1 adopts an open conformation (left), whereas the 48S complex without 5MP1 is predominantly closed (right). In the 5MP1-bound structure, eIF2 (blue), initiator tRNA^Met^ (green), and 5MP1 (purple) are shown as cartoons. Initiator tRNA^Met^ in the 48S complex without 5MP1 is shown in dark green. **(B)** In the presence of 5MP1, the initiator tRNA^Met^ adopts a Pout conformation (left), 7 Å away (arrow) from the Pin conformation in the closed 48S complex without 5MP1. The mRNA in the 5MP1-bound complex is shown in pink, and in the 48S complex without 5MP1 in light pink. **(C)** The close-up shows no codon-anticodon pairing in the 5MP1-bound complex, in contrast to the well-resolved anticodon-paired AUG in the 48S complex without 5MP1. **(D)** An overlay of the 5MP1 model with the 48S closed-head conformation (superposition) shows a steric clash between the 5MP1 N-terminal tail and the helix 18 of 18S rRNA. Experimental 5MP1 model is shown in purple and superimposed 5MP1 in lilac.

5MPs consists of two domains, which contact eIF2, initiator tRNA^Met^, eIF1A and the mRNA tunnel (Fig. 5A). The C-terminal domain (residues 248-407) docks onto eIF2β and initiator tRNA^Met^. Here, Tyr351 and Ile354 of the W2/HEAT3 domain—previously implicated in interactions with eIF2β (*23, 26*)—bind the universally conserved His230 of eIF2β through π-π stacking (Tyr351) and hydrophobic packing (Ile354) (Fig. 5B). A Tyr351Ala/Ile354Ala double mutant (Y351A/I354A) of 5MP1 renders the protein ∼4-fold less inhibitory than wild-type 5MP1 on a transcript containing a CUG start codon in a strong Kozak context (Fig. 5F; fig. S9), highlighting the importance of this contact for 5MP function. Residues Phe401/Val402/Trp404/Leu405 of 5MP1, identified as part of the 7A mutation and implicated in the interaction with the eIF2β N-terminus (*23*), are resolved in our map, and nearby densities may correspond to eIF2β N-terminus, however, unambiguous assignment is not possible due to insufficient resolution, leaving open the possibility of transient interactions (Fig. 3; fig. S7C; fig. S10). This possibility is consistent with suggestions that the intrinsically disordered N-terminal tail of eIF2β engages 5MP through dynamic and competitive contacts at its lysine-rich boxes (*26, 59*). Residues in the previously characterized BN1 mutant (Lys380/Lys383/His386/Lys391) (*23*) are located near eIF2β (fig. S10), and the poorly resolved density noted above for 5MP1-7A and eIF2β also lies in proximity to these residues. Alternatively, the unresolved density may correspond to the N-terminal segment of eIF3c rather than the eIF2β N-terminal tail, as the former was shown to stabilize 5MP1 in the 48S complex (*23*). Given the ambiguity and low resolution of this density, we also cannot exclude that the density reflects noise rather than one of these interactions. Flexible low-occupancy interactions are challenging to resolve by cryo-EM. Crosslinking coupled to mass spectrometry or other approaches would help corroborate these observations and map the contact residues.

**Fig. 5.**
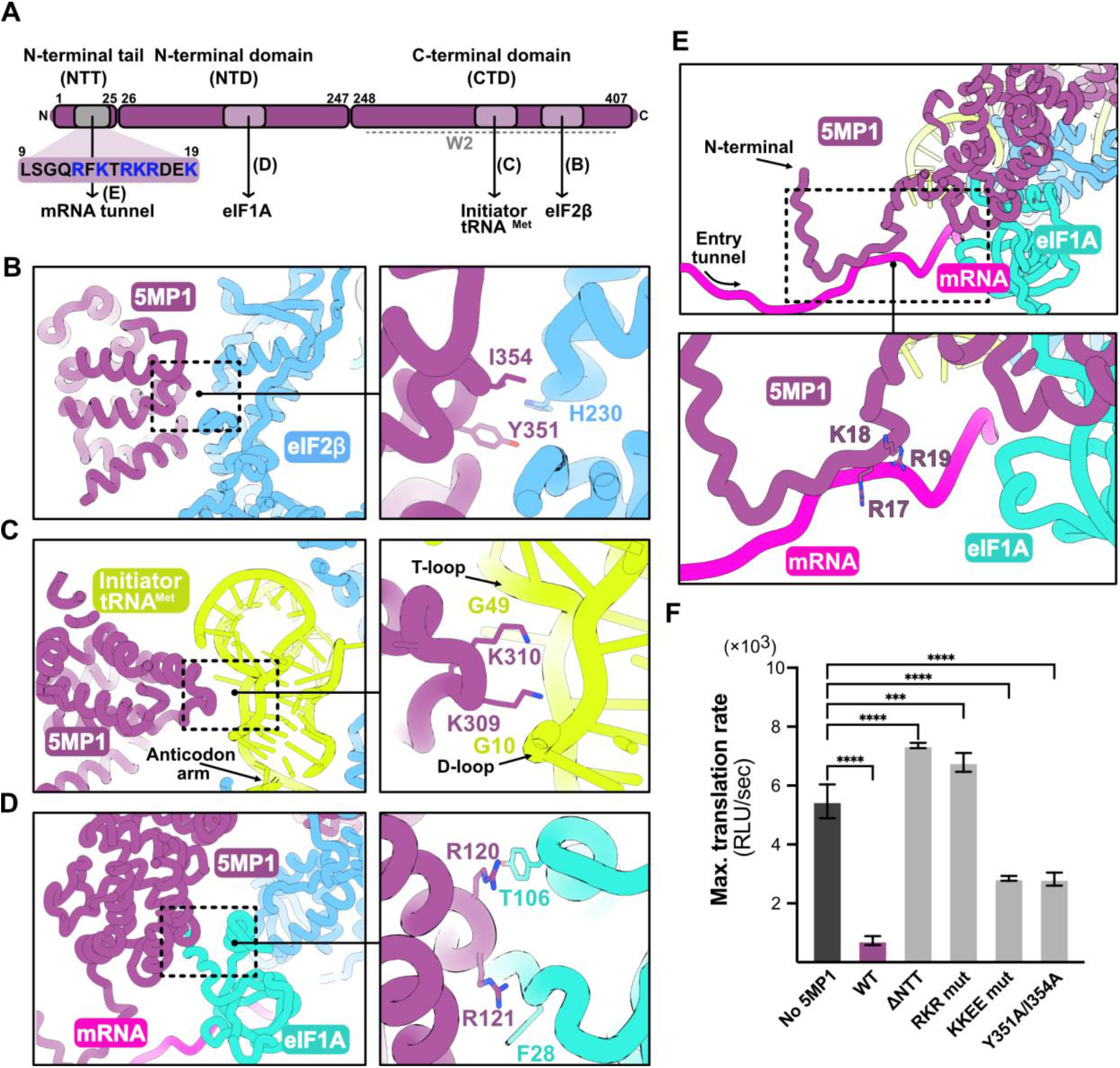
Interactions of 5MP1 within the 48S pre-initiation complex. **(A)** Schematic representation of 5MP1 domain organization, interactions within the 48S complex and the conserved sequence in the N-terminal tail (NTT). **(B)** Close-up view of the 5MP1-eIF2β, **(C)** 5MP1-initiator tRNA^Met^ and **(D)** 5MP1-eIF1A interfaces, highlighting the interacting residues. **(E)** Close-up view of NTT highlights the Arg-Lys-Arg motif facing the mRNA backbone (right). **(F)** Effects of structure-guided mutations on 5MP1-mediated inhibition of translation from a transcript containing a CUG start codon in a strong Kozak context, measured at 1 μM (mean ± SEM, n=3): wild-type 5MP1; the NTT-deletion mutant (ΔNTT; residues 1-25); the RKR(17-19)-to-AAA mutant (RKR mut); the KKEE(309-312)-to-EEQQ mutant (KKEE mut); and the Y351A/I354A double mutant. Data represent mean ± SD (n=3). Statistical significance was assessed by one-way ANOVA; ****P < 0.0001, ***P < 0.0002, **P < 0.0021, ns = not significant. 5MP1 is shown in purple, eIF2 in blue, eIF1A in turquoise, and the initiator tRNA^Met^ in green.

Residues Lys309 and Lys310 of 5MP1 reach toward the phosphate groups of D-loop and variable-region nucleotides 10 and 49 of initiator tRNA^Met^, in the vicinity of eIF2β (Fig. 5C). This interaction appears as important for 5MP1 function as the contact with eIF2β described above. Specifically, substituting the Lys309/Lys310/Glu311/Glu312 (KKEE) motif with Glu/Glu/Gln/Gln (EEQQ) to neutralize this positively charged surface reduced 5MP1-mediated inhibition by nearly 4-fold on the same CUG reporter (Fig. 5F; fig. S10; KKEE mut). Because eIF5 shares a homologous C-terminal W2/HEAT3 domain and binds eIF2β (*22, 43*), we asked whether the eIF5 CTD would similarly interact with the 48S scanning-competent complex and whether the contacts with eIF2β and initiator tRNA^Met^ reported here are compatible. Superposition of eIF5 CTD onto 5MP1 CTD within the 48S•5MP1 complex suggests similar interactions with the 48S complex (fig. S11A), consistent with their competition at this site (*60*). However, despite the overall similarity, there are differences between 5MP CTD and eIF5 CTD interactions within the 48S: (*1*) eIF5 residues 366-368 can make additional contacts with eIF2β that are not observed with 5MP1 and (*2*) the eIF5 helix reaching toward initiator tRNA^Met^ is shorter compared to the one in 5MP and is likely incompatible with interactions with the initiator tRNA^Met^ (fig. S11B). These differences highlight that the CTD of 5MP1 and eIF5 engage the 48S through distinct molecular interactions despite their shared domain architecture. Additionally, 5MP1 NTD interacts with the 48S complexes (see discussion below), which may be essential for stabilizing this interaction, consistent with the inability to resolve eIF5 CTD in numerous 48S complexes that have a clearly resolved eIF5 NTD (*26*).

Lastly, the 40S A site accommodates the 5MP1 N-terminal domain (NTD) next to eIF1A, stabilized by cation-π interactions between 5MP1 (Arg120 and Arg121) and eIF1A (Tyr106 and Phe28; Fig. 5D). The N-terminal tail (NTT) extends into the mRNA tunnel (Fig. 5E), where the terminal residues form a β-stand with protein uS3 and the conserved Arg17-Lys18-Arg19 motif may be involved in contacts with the mRNA backbone. Transitioning to the closed-head codon-recognition state would require displacement of NTT to prevent the positively charged Arg-Lys-Arg motif of NTT from clashing with helix 18 of 18S rRNA (Fig. 4D) (*55*). This steric hindrance is consistent with the absence of 5MP1 in the closed-head maps. To test the functional significance of the NTT, we generated two mutants. Deleting the entire NTT (ΔNTT; residues 1-25) or mutating the Arg17/Lys18/Arg19 motif to three alanines alone (RKR mut) abolished 5MP1’s inhibitory activity on a transcript containing a CUG start codon in a strong Kozak context, relative to wild-type 5MP1 (Fig. 5F; fig. S10). The comparable magnitudes of these effects suggest that the RKR motif alone accounts for the functional contribution of the NTT to 5MP1-mediated inhibition. Indeed, the RKR motif is among the most highly conserved elements across 5MP homologs, from early-diverging unicellular lineages to vertebrates (Methods; fig. S12), emphasizing that NTT-mediated engagement in the mRNA tunnel represents an ancient and conserved mechanism for controlling start-codon fidelity.

Taken together, these findings reveal that 5MP1 interacts with the 48S PIC through multiple contacts—spanning the NTT, NTD, and CTD—that collectively bias the complex toward the open-head, scanning-competent state. The residues directly mediating 5MP1’s contacts within the 48S PIC are fully conserved in 5MP2 (fig. S9D), indicating that 5MP2 is poised to engage the 48S PIC through the same mechanism. The most pronounced sequence divergence is located in a short segment (residues ∼88-93) immediately following the NTD (fig. S10). Superposition of an AlphaFold2-predicted structure of 5MP2 suggests that this region may form a flexible loop near the 18S rRNA of the 40S beak (fig. S10B) that may influence 5MP2 binding or interaction of the NTT with the mRNA tunnel, in keeping with the higher IC50 for 5MP2 (Fig. 1C-D). Post-translational modifications represent another layer of functional regulation: both 5MP paralogs contain a conserved Casein Kinase 2 (CK2) phosphorylation site within their CTD, homologous to a site in eIF5 whose phosphorylation modulates eIF2β binding (*26*). These functional consequences remain to be explored in future work. Together, the unique structural contacts emphasize that 5MP1 function extends beyond being just an eIF5 competitor and define it as a *bona fide* initiation factor that stabilizes the scanning-competent 48S state.

### The mechanism of initiation regulation by 5MPs

The molecular mechanism of eIF5-mimic proteins has long remained unknown, despite their key roles in regulating translation initiation and in diseases such as cancer. Our work shows that 5MP potently enforces start-codon fidelity by inhibiting initiation at alternative start sites, such as non-AUG codons or AUG in suboptimal Kozak contexts (Fig. 1). In the c-Myc context, this regulation shifts translation from a mixture of AUG- and non-AUG-initiated proteoforms toward predominantly the AUG-initiated form, while also enhancing overall translation, a dual effect with potential relevance in disease (Fig. 2). The cellular concentrations of 5MP1s are regulated by the weak start codons in 5MP-encoding genes, offering a surveillance mechanism for accurate selection of AUG codons and switching between proteoforms (*24*).

To further delineate 5MP function in the context of cellular components, we performed *in extracto* cryo-EM, achieving near-atomic resolution for native 5MP-bound initiation complexes (Fig. 3). Cryo-EM analyses reveal that 5MP binds exclusively within the open-head 48S complexes, suggesting a straightforward structural mechanism of 5MP action (Fig. 4; Fig. 6). To reduce initiation at a suboptimal start site, 5MP shifts the equilibrium between 48S conformations toward a widely open, scanning-competent state. Within 48S, 5MP interacts with eIF2β, initiator tRNA^Met^, eIF1A, and the mRNA tunnel (Fig. 5). The NTT of 5MP is folded within the mRNA entry tunnel widened by the head opening (Fig. 5), and head closure is incompatible with this position due to a steric hindrance (Fig. 4D), highlighting the NTT as the functional core of 5MPs. The transition of 48S into the closed codon-recognition state (*e.g.*, on a strong AUG codon) requires eIF2β displacement (*7*), dissociation of eIF1, and the closure of the 40S head. This must coincide with or succeed 5MP dissociation, as is indicated by the absence of 5MP in closed 48S states and by the incompatibility of the NTT with the closed 48S. One possibility is that 5MPs associate with the scanning 48S reversibly, and that its binding is prevented when a strong codon recognition biases fast head closure (*7, 55*). Alternatively, for a stably bound 48S•5MP scanning complex, an early interaction of the eIF2•initiator tRNA^Met^ and/or other initiation factors with the AUG codon and Kozak context may induce a transient conformational rearrangement that reduces the affinity of 5MP for the 48S. This is consistent with prior work showing that eIF1 can dissociate from the preinitiation complex upon codon-anticodon pairing at a strong AUG codon (*61*). These scenarios may explain why 5MPs have little effect on strong AUG/Kozak contexts, where codon recognition outcompetes 5MP binding, while they strongly inhibit initiation at weak or non-AUG codons subject to a slower initiation (Fig. 1B).

**Fig. 6.**
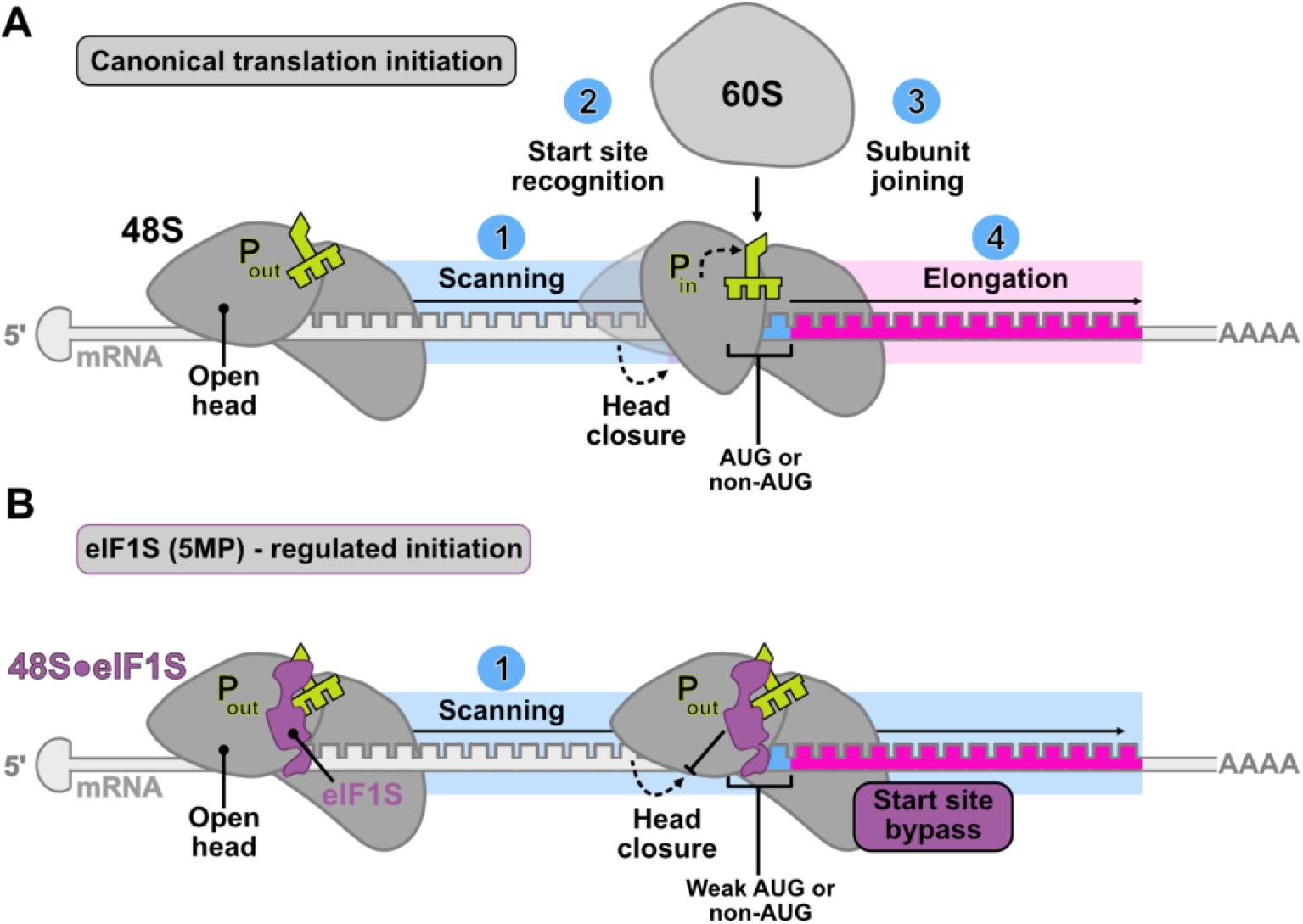
Schematic of eIF1S (5MP)-mediated regulation of start codon recognition. **(A)** During canonical initiation, the 48S complex scans the mRNA until it recognizes a start codon. Both canonical AUGs and suboptimal codons, including weak AUGs and non-AUGs, can trigger scanning arrest. This arrest is accompanied by a conformational shift of the 48S head from an open to a closed state, while the initiator tRNA^Met^ transitions from the Pout to the Pin position to allow codon-anticodon pairing. Successful start-site recognition then drives 60S joining and elongation. **(B)** By contrast, eIF1S (5MP) stabilizes the 48S complex in a scanning-competent state, maintaining the 48S head in the open conformation and the initiator tRNA^Met^ in the Pout position. As a result, weak AUG and non-AUG codons fail to induce arrest, are bypassed during scanning, and do not initiate elongation

Together, our findings elucidate how 5MPs add a robust layer of stringency to canonical initiation factors and shift our understanding of their structural roles in translation. Rather than acting solely through competition with eIF5, 5MP functions as a *bona fide* initiation factor that binds the 48S complex and biases it toward scanning, preventing commitment at suboptimal start codons. To reflect the direct roles of 5MPs within the 48S pre-initiation complex, we propose designating them as eIF1S (eukaryotic initiation factor 1 scanning; eIF1S-1 for 5MP1 and eIF1S2 for 5MP2). This mechanistic and structural insight into 5MP-mediated proteoform regulation may lead to therapeutic developments targeting diseases driven by or dependent on dysregulated initiation.

## Acknowledgments

We thank Chen Xu, Kangkang Song, and Christna Ouch for support with cryo-EM data collection at UMass Chan Medical School; Johannes Elferich, Dongjie Zhu, and Stephen Diggs for valuable scientific input; and members of the Grigorieff and Korostelev laboratories for constructive feedback on this study. We also thank Assen Marintev and Thomas Hainthaler for helpful discussions on datasets.

## Author contributions

**X.Z.** performed and analyzed biochemical experiments, prepared cell cultures and cryo-EM samples; collected and analyzed cryo-EM data; prepared illustrations and drafted the manuscript. **C-Y.H.** performed and analyzed biochemical experiments; prepared cryo-EM samples; prepared illustrations and drafted manuscript. **Z.S.** developed cell lysate protocols. **N.G.** designed research, supervised data analysis, drafted manuscript and secured funding. **A.A.K.** designed research, supervised data analysis, drafted the manuscript and secured funding. All authors revised the manuscript.

## Data, code, and materials availability

All data and code needed to evaluate and reproduce the results in the paper are present in the paper and/or the Supplementary Materials.

## Materials and Methods

### BSC-1 cell culture and cell lysate preparation

BSC-1 cells (ATCC), a well-established epithelial cell line historically central to virology and membrane trafficking research, were cultured in DMEM (Invitrogen) supplemented with 10% heat-inactivated FBS, 2 mM GlutaMAX (Gibco) and 1 % (v/v) of 100× Penicillin-Streptomycin (Gibco) (*62*). Cells were seeded in a 75 cm² flask at a density of ∼20,000 cells/flask and cultured to confluence. After being washed twice with pre-warmed PBS, the cells were detached with Trypsin-EDTA (Gibco) and collected by centrifugation at 300×g for 4 minutes. The cell pellet was resuspended in 100 μL of semi-permeabilization buffer (25 mM HEPES, 110 mM KOAc, 15 mM Mg(OAc)_2_, 1 mM DTT, 0.015% digitonin, 2× protease inhibitor cocktail (Roche), 20U SUPERase-In RNase inhibitor (Thermo Fisher Scientific: AM2696), 1 mM EGTA) at 4 °C for 5 min. Following centrifugation at 1,000×g for 5 minutes, the supernatant was collected for grid preparation (see below). RNA concentration was quantified before grid preparation as a quality control measure using a NanoDrop One Spectrophotometer (A260/A280 and A260/A230; Thermo Fisher Scientific) without diluting the extract.

### 5MP protein expression and purification

Coding sequences for human 5MP1 (UniProt:Q9Y6E2), 5MP2 (UniProt: Q7L1Q6), 5MP1 ΔNTT mutant (deletion of residues 1-25 and mutation of Pro26Ala), 5MP1 RKR mutant (RKR(17–19)-to-AAA), 5MP1 KKEE mutant (KKEE(309-312)-to-EEQQ) and Y351A/I354A double mutant; were synthesized (GenScript) and cloned into pET29a(+) using NdeI-XhoI restriction sites, generating constructs with a C-terminal His_6_ tag (GenScript). All plasmids were sequence-verified before use. Plasmids were transformed into *E. coli* BL21(DE3) and cultures were grown in LB medium at 37 °C to an OD600 of 0.6, induced with 0.5 mM IPTG, and expressed at room temperature for 5 hours. Cells were harvested by centrifugation and stored at −80 °C until purification. The cell pellets were resuspended in a lysis buffer (20 mM Tris-HCl pH 8.0, 500 mM NaCl, 5 mM imidazole, 0.5 mM DTT, 2× protease inhibitor cocktail (Roche),) and lysed by sonication on ice. Lysates were clarified by centrifugation at ∼40,000×g for 40 min at 4 °C, and supernatants were filtered through a 0.22 μm filter before chromatography. The clarified lysate was loaded onto a pre-equilibrated Ni-NTA affinity column, washed with 10 column volumes of binding buffer (20 mM Tris-HCl pH 8.0, 500 mM NaCl, 5 mM imidazole, 0.5 mM DTT), and eluted on an ÄKTA FPLC (Cytiva) system using a linear gradient up to 1 M imidazole. Fractions were analyzed by SDS-PAGE, and peak fractions pooled. The pooled Ni-NTA eluate was dialyzed overnight against a storage buffer (20 mM Tris-HCl, pH 7.8, 125 mM NaCl, 0.5 mM DTT), concentrated using 10 kDa Amicon filters and concentration estimated using UV absorbance at 280 nm on a NanoDrop One Spectrophotometer (Thermo Fisher Scientific). A total of 100 mg of protein was loaded into a HiTrap Q HP anion-exchange column pre-equilibrated (20 mM Tris-HCl, pH 7.8, pH 8.0, 125 mM NaCl, 0.5 mM DTT). Proteins were eluted with a linear NaCl gradient. Fractions containing single bands corresponding to 5MP1 or 5MP2 by SDS-PAGE were pooled, dialyzed overnight against a storage buffer (10 mM HEPES pH 7.5, 125 mM NaCl, 0.5 mM DTT), concentrated using 10kDa Amicon filters and concentration estimated using UV absorbance at 280 nm on a NanoDrop One Spectrophotometer (Thermo Fisher Scientific). Proteins were subjected to a final purification using Superdex 200 Increase 10/300 GL column equilibrated in 10 mM HEPES pH 7.5, 125 mM NaCl, 1 mM TCEP. For 5MP1, fractions corresponding to elution peaks 6-9 were collected; for 5MP2, fractions 3-6 were recovered. The final preparations showed a major peak migrating at ∼42 kDa. Protein concentration was determined by UV absorbance at 280 nm using theoretical extinction coefficients and a NanoDrop One Spectrophotometer (Thermo Fisher Scientific), with purity confirmed by SDS-PAGE (NuPage Bis-Tris Mini Protein Gels 4-12%, MOPS SDS buffer kit, SeeBlue Pre-stained protein standard (Invitrogen)). Aliquots were flash-frozen in liquid nitrogen and stored at −80 °C until use. All the buffers used were filtered with 0.5 μm filters.

### Plasmids for mRNA preparations

Plasmids used for this mRNA preparation are listed in Table S3. A plasmid carrying the coding sequence for NanoLuciferase (NLuc) flanked by the 5′ and 3′ UTRs of rabbit β-globin was synthesized by Azenta (vector: pUC-GW-Kan), as designed and described (*63*). Variants of the plasmid containing the mutations in the AUG or CUG start codon, and their Kozak contexts, were also synthesized by Azenta (vector: pUC-GW-Kan).

The plasmids containing the 5′UTR human *c-MYC* (NM_002467.6) with in-frame start codons (CUG and AUG) and the coding sequence for the first 48 nucleotides of c-MYC, fused to a 3×FLAG and NLuc CDS was synthesized by Azenta (vector: pUC-GW-Kan). Variants of the plasmid containing the mutations in the start codons and their Kozak contexts were also synthesized by Azenta (vector: pUC-GW-Kan). The constructs are listed in Table S3. All plasmids were sequenced to verify the sequences.

### *In vitro* transcription of NanoLuciferase reporter mRNAs

DNA templates for *in vitro* transcription of mRNAs were PCR-amplified from the plasmids using primers flanking the template region for transcription. The T7 promoter sequence and a 30-nt poly(A) tail were added through overhangs of the forward and reverse primers. The following forward primer sequences were used: *TTTTTTAATACGACTCACTATAGGGAGA*ACACTTGCTTTTGACACAACTGTG (NLuc constructs, 5′-3′); *TTTTTTAATACGACTCACTATAGGGAGA*AACTCGCTGTAGTAATTCCAGCGAG (c-Myc constructs, 5′-3′); sequence with the T7 promoter site is in italic.

The reverse primer, including a poly(A)-encoding sequence in all constructs, was (5′-3′): TTTTTTTTTTTTTTTTTTTTTTTTTTTTTTGCAATGAAAATAAATTTCCTTTATTAGCC.

PCR was performed using Phusion High-Fidelity DNA Polymerase (NEB; M0530L) according to NEB protocol. After PCR, DNA templates were purified by phenol/chloroform extraction and dissolved in nuclease-free Milli-Q water. DNA concentrations were measured using a NanoDrop One Spectrophotometer (Thermo Fisher Scientific). The presence of a single band as a DNA template was confirmed by agarose gel electrophoresis (1% (w/v) agarose in TAE buffer). *In vitro* transcription reactions were carried out using 4 μg of purified DNA templates and purified recombinant T7 polymerase in transcription buffer (166 mM HEPES-KOH, pH 7.5; 20 mM MgCl_2_; 40 mM DTT, 2 mM spermidine, 25 mM each of ATP, GTP, CTP, and UTP; and 40 U/μl RNase Inhibitor (NEB; M0314S)) in an 80 μl reaction. After incubation at 37 °C for 3.5 hours, magnesium pyrophosphate precipitate was removed by centrifugation (14,000 x g, 5 min), and mRNA was precipitated from the supernatant by adding 7.5 M LiCl (2.5 M final concentration) and incubating at −80 °C overnight. The next day, mRNA was pelleted by centrifugation (21,300 x g, 15 min at 4 °C), washed with cold 80% ethanol and pelleted again. This washing step was repeated three times. After discarding the supernatant, the mRNA pellet was air-dried and dissolved in nuclease-free Milli-Q water. To attach 5′cap, capping reactions were performed using the Vaccinia Capping System (NEB; M2080S). The 5′capped mRNA was then purified by LiCl precipitation as described above and dissolved in nucleases-free Milli-Q water. mRNA concentration was determined from UV absorbance at 260 nm using a NanoDrop One Spectrophotometer (Thermo Scientific). The size and integrity of the *in vitro* transcribed mRNA were examined by denaturing agarose gel electrophoresis (1% (w/v) agarose in MOPS buffer with 1.11% (v/v) formaldehyde) alongside an ssRNA ladder (NEB). The stock solution of *in vitro* transcribed mRNAs was stored at −80 °C.

### *In vitro* mRNA translation in RRL

*In vitro* translation was performed using a commercial micrococcal-nuclease-treated RRL (Promega; L4960) with modifications as described below. Translation reactions were carried out in the presence of 50% RRL, 30 mM HEPES-KOH (pH 7.5), 50 mM KOAc, 1.0 mM Mg(OAc)_2_, 0.2 mM ATP, 0.2 mM GTP, 0.04 mM of 20 amino acids (Promega), 5 mM DTT, and 1% furimazine NanoLuciferase substrate (Promega; N113A). To initiate translation, 11 µL of reactions were preincubated at 30 °C for 3 min, followed by addition of *in vitro* transcribed mRNAs encoding Nanoluciferase (10 nM final) to a total volume of 12 µL. Translation kinetics were monitored by continuously recording NanoLuciferase luminescence at 30 °C for 20 min using an Infinite m1000 pro microplate reader (Tecan). The luminescence values were plotted over time using Prism 10 (GraphPad Software). Progress curves were recorded for experiments in triplicates. The maximum translation rates were calculated from the maximum rate of luminescence change (Max ΔRLU/Δsec), which were obtained as the peak values of the first derivative of the RLU curves calculated in Prism 10 (GraphPad Software).

### Determination of IC50 of 5MP1/2

To derive the IC_50_ values for 5MP1/2-mediated translation inhibition from luminescence curves, maximum translation rates (Max ΔRLU/Δsec) were plotted as a function of 5MP1/2 concentrations. The IC_50_ values were determined by fitting the data using a four-parameter inhibition model (Y=Bottom + (Top-Bottom)/[1+(IC_50_/X)^HillSlope]) with nonlinear regression (curve fitting) in Prism 10. The average translation rate in the absence of 5MP1/2 was defined as 0% inhibition (maximum translation activity), whereas the average rate at the highest 5MP1/2 concentration was set as 100% inhibition (minimum translation activity).

### Detection of RNase contamination within recombinant 5MP1/2 purified protein solutions

To test for RNase contamination in the purified recombinant 5MP1/2 proteins, NLuc mRNAs (final 140 nM) were incubated with and without the purified recombinant 5MP1/2 (final 1 µM) at 37 °C for 1 hour, *i.e.,* exceeding the extents of translation experiments in this work. As a control, incubation reactions were also set up with 40 units (U) of RNase Inhibitor (NEB; M0314S) for a final volume of 20 μl. The reactions were stopped by the addition of 2× RNA loading dye (NEB; B0363S), and the samples were heated at 70 °C for 10 min. mRNA integrity was assessed by 1% denaturing formaldehyde agarose gel pre-stained with SYBR Safe DNA gel stain (Invitrogen; S33102).

### Western blotting

Equal volumes of RRL translation reactions were mixed with 2× urea sample dye (80 mM Tris-HCl, pH 6.8; 138.75 mM SDS; 8 M urea; 3 mM bromophenol blue; 16% (v/v) glycerol) and boiled at 98 °C for 10 min. The total proteins in RRL samples were separated on 4-20% Mini-Protean TGX Precast Protein gels (Bio-Rad) alongside a protein marker (Bio-Rad) at 130 V for 2.5 h in 1× Tris-Tricine-SDS running buffer, and proteins were transferred to a PVDF membrane using a Trans-Blot Turbo transfer system (Bio-Rad).

To reduce non-specific signal, the membranes were blocked in blocking buffer (1× PBST containing 0.1% Tween-20 and 5% (v/v) non-fat dry milk (Research Products International; M17200)) at room temperature for 30 min. The membranes were then incubated either with 1:3000 mouse anti-FLAG M2 IgG1 monoclonal antibody (Sigma-Aldrich; F1804) or 1:5000 rabbit anti-BZW2 IgG polyclonal antibody (GeneTex; GTX106985) in blocking buffer at 4 °C overnight. After washing the membranes by shaking in 1× PBST for 10 min three times, the membranes were incubated either with 1:6000 goat anti-mouse IgG secondary antibody conjugated to horseradish peroxidase (Invitrogen) or with 1:10000 goat anti-rabbit IgG secondary antibody conjugated to horseradish peroxidase (Invitrogen) in blocking buffer at room temperature for 1 h. After washing the membranes three times, 10 min each, the blots were detected using SuperSignal West Atto Ultimate Sensitivity Substrate (Thermo Fisher) and imaged on the ChemiDoc MP system (Bio-Rad). The intensities of Western blot bands were quantified using ImageJ.

### RRL sample preparation for cryo-EM

Purified 5MP1 (5 μM final) was added to 60 % commercial micrococcal-nuclease-treated RRL (Promega; L4960) supplemented with 30 mM HEPES-KOH (pH 7.5), 50 mM KOAc, 1.0 mM Mg(OAc)_2_, 0.2 mM ATP, 0.6 mM Guanosine 5′-[β,γ-imido]triphosphate (GMPPNP) (*47–51*), 0.04 mM of 20 amino acids (Promega) and 20U SUPERase-In RNase inhibitor (Thermo Fisher Scientific: AM2696), maintained on ice until mRNA addition. Using GMPPNP in the RRL allowed us to capture ∼3 times more inititiation complexes than in the endogenous BSC-1 lysate. NLuc (AUG within a moderate Kozak sequence) was added to the 5MP1-RRL-supplemented mixture and incubated for 10 min at 30 °C under continuous agitation (400 rpm). The mixture was directly added to the grids and vitrified.

### Grid preparation and plunge freezing

For BSC-1 cellular lysates, Quantifoil Au 300 mesh R2/2 holey SiO₂ grids (Electron Microscopy Sciences) were glow-discharged in an EMITECH K100X (25 mA, negative polarity, 45 s). Grids were back-blotted for 8 s and plunge-frozen in liquid ethane at -184 °C using a Leica EM GP (15 °C, 85% relative humidity). Vitrified grids were stored in liquid nitrogen until imaging.

For 5MP1-RRL samples, Quantifoil R2/1 holey-carbon grids coated with a thin continuous carbon layer (Electron Microscopy Sciences) were glow-discharged in a PELCO easiGlow (20 mA, negative polarity, 30 s). A Vitrobot Mark IV (Thermo Fisher Scientific) was equilibrated to 4 °C and 100% relative humidity, with blot force set to zero. For each sample, 2.5 µL of lysate was applied, immediately blotted (2-3 blots, 3-7 s each, force 0), and plunge-frozen in liquid nitrogen-cooled ethane. Grids were stored in liquid nitrogen.

### Cryo-EM data collection and analysis

Datasets for BSC-1 cell lysate and 5MP1-RRL samples were collected on a Titan Krios electron microscope (Thermo Fisher Scientific) operating at 300 kV, equipped with a Gatan Imaging Filter (20-eV slit width) and a K3 direct electron detector (Gatan). Images were recorded at a nominal magnification corresponding to a super-resolution pixel size of 0.415 Å (physical pixel size 0.83 Å), using a target defocus range of 1.0-1.5 μm. Automated data acquisition was performed in SerialEM (*64, 65*) using beam-image shift to collect 5 movies per hole across 9 holes at each stage position. Zero-loss peak (ZLP) refinement was performed every 90 mins at a predetermined location to avoid darkened areas. The BSC-1 dataset consisted of 162,583 movies and the 5MP1-RRL dataset of 104,673 movies; each movie containing 30 frames with an exposure of ∼1 e⁻ Å⁻² per frame (total exposure: ∼30 e⁻ Å⁻²). Beam-induced motion correction, gain reference application, exposure weighting, and 2×/4× binning (to pixel sizes of 0.83 Å and 1.66 Å, respectively) were performed using MotionCor2(*66*). CTF parameters were estimated using CTFFIND (*67*) via the *cis*TEM GUI (*68*). For 2D template matching (2DTM), atomic coordinates for the 40S body from PDB 7R4X were converted into a 3D map using the simulate program in *cis*TEM (*69*), with a pixel size of 1.66 Å, 336×336×336 box, and a linear B-factor of 10. 2DTM was executed on 4× binned images (pixel size 1.66 Å), processing data in blocks of 2,000 images. This yielded 523,233 targets for BSC-1 lysate and 1,573,843 targets for 5MP1-RRL, using in-plane angular sampling of 1.5°, out-of-plane sampling of 2.5°, and no defocus search. Template-matched coordinates, Euler angles, and CTFFIND5-derived defocus values were exported using the MT package in *cis*TEM GUI to generate .star files, which were merged using an in-house Python script; paths to the 4× binned images were replaced with those for 2× binned images (pixel size 0.83 Å). The updated .star file was re-imported in the MT package, and particle stacks (box size 560³ pixels) were generated using the *cis*TEM refine pipeline. Initial 3D reconstructions were computed using Generate 3D, and final alignment parameters and defocus values were exported to Frealign format (fig. S5). Collection and processing pixel sizes were as follows: super-resolution, 0.415 Å; physical, 0.83 Å (final reconstructions); 2× bin, 1.66 Å (CTF estimation and 2DTM); 4× bin, 3.32 Å (3D classification); and 8× bin, 6.64 Å (3D classification). Empiar ID-13853 and 13577 (BSC-1 cell lysate) and Empiar ID 13852 (RRL).

### Cryo-EM data classification

All classifications steps were performed in Frealign v.9.11 (*70*) on binned data (8×, 4×, 2×), with particle stacks prepared using the program resample in *cis*TEM package (*68*). Binning of the particle stacks was performed using the program resample from cisTEM. Initial unmasked 3D maximum-likelihood classification separated elongating ribosomes (80S) and junk from pre-initiation complexes (8× binned data). For BSC-1 cell lysates, pre-initiation complex (PIC) classes were merged using an in-house Python script, preserving micrograph paths and generating a .star file compatible with *cisTEM* (available in YAFW: https://github.com/GrigorieffLab/yafw). A particle stack of PIC was generated in *cis*TEM, and further manual refinements were performed, including cycles of x,y alignment, Euler angles, and defocus/beam-tilt refinement. Focus-masked 3D maximum-likelihood subclassification was then applied to the head region (mask diameter 90 Å) to separate subpopulations. Classes containing the unknown density at the A site were merged using YAFW, and 3D reconstructions were performed from unbinned data (pixel size 0.83 Å) in *cis*TEM. Local resolution estimation and map denoising were performed using RELION-3.1 (*71*).

For 5MP1-RRL samples, an initial classification into 100 classes was performed. Representative PIC classes were divided into six complex types and merged using YAFW scripts. Particle alignment files (.star) were used to generate stacks in *cis*TEM for further refinement, as described above. Focus-masked subclassification focused on 5MP, the head, eIF2, or mRNA/initiator tRNA^Met^ codon-anticodon sites did not reveal additional heterogeneity, even when tested on 2×, 4×-, and 8× binned data. The 2DTM and classification workflow are illustrated in fig. S5A. 3D reconstructions with unbinned data (pixel size 0.83 Å) were performed in *cis*TEM. Local resolution estimation and map denoising were performed using RELION-3.1. Maps are deposited in EMDB, EMD-78242 (48S•5MP1), EMD-78324 (48S closed head), EMD-78325 (48S-like•5MP1).

### Protein identification in a cryo-EM map

The map containing the unknown density was low-pass filtered (4 Å) and segmented, retaining only the density corresponding to the unknown feature. ModelAngelo (*36*) implemented in RELION-5.0 (*72*) was run without the sequence module (no_seq option) to generate an initial model based on HMM profiles. These profiles were then searched against the reference proteome using the hmm_search option. Hits for 5MP and 5MP2 showed the highest significance, with E-values of 1.43 × 10⁻⁹ and 1.24 × 10⁻³⁷, respectively. The predicted AlphaFold2 structures of these proteins (AF-Q9Y6E2) were fitted into the density to validate the identification(*37, 73*).

### Model building

Maps were sharpened using DeepEMhancer (*74*). The initial models were derived from the *Oryctolagus cuniculus* 80S ribosome (PDB 9BDL) (*75*), the *H. sapiens* 48S complex with eIF4F/eIF4A (PDB 8OZ0)(*4*), and the *H. sapiens* 48S complex in the open scanning state (PDB 8PJ1) (*7*). The 18S rRNA from 9BDL was first rigid-body-fitted to the density. Then, ribosomal proteins and initiation factors from 8OZ0 or 8PJ1 were independently rigid-body-fitted using ChimeraX (*76*). The 5MP1 model was generated using AlphaFold2 prediction (AF-Q9Y6E2) was rigid-body fitted into the density. Following initial placement, all components were refined using ISOLDE (*77, 78*) within ChimeraX, applying segment-wise interactive molecular dynamics. Final refinements were performed with phenix.real_space_refine, yielding models with good stereochemical quality and high correlation with the cryo-EM maps (Table S1). The model-map FSC was calculated using Phenix (*79*). PDB identifier 37KM.

### Head rotation measurements

To compare head rotation between initiation complexes, we generated a closed-head model by rigid-body fitting the human 48S complex in the AUG-recognition state (PDB 8PJ2) into our 2.8-Å map using ChimeraX (*76*), followed by a single round of real-space refinement in Phenix (*78, 79*). For both the 5MP1•48S (open-head) and the refined closed-head model, we aligned the 18S body (nucleotides 1-1213 and 1691-1869) using Matchmaker in ChimeraX and saved the aligned coordinates. The 18S head (nucleotides 1214-1690) was then independently aligned, and the resulting angular displacement was calculated in ChimeraX. Head rotation between our 48S•5MP1 complex and the previously reported 48S open-head structure (PDB 8PJ1) was measured using the same procedure (*7*).

### Multiple sequence alignment

Protein sequences used for comparative analysis were obtained from a previously published phylogenetic dataset (*22*) and supplemented with mammalian organisms with their accession numbers as follows:

*Homo Sapiens (UniProt:Q9Y6E2); Giardia intestinalis* (*UniProt:*C6LQG9); *Chlamydomonas reinhardtii (RefSeq:*XP_001691456, *UniProt:*A8IP37); *Volvox carteri (UniProt:*D8TS79); *Chlorella variabilis (UniProt:*E1Z485); *Coccomyxa subellipsoidea (UniProt:*I0Z2Z0), *Selaginella moellendorffii (UniParc:*UPI0001E02D4B); *Physcomitrella patens* (*UniParc*: UPI000162096B); *Ricinus communis (UniParc:*UPI0001940808); *Arabidopsis thaliana* (*UniParc*:UPI00000A2E64); *Triticum aestivum (UniParc:*UPI00029B5BFD); *Zea mays* (*UniParc*:UPI00017B6F18); *Chondrus crispus (UniParc:*UPI000331C39F); *Schizophyllum commune* (*UniParc*: UPI0001DF4DBE); *Monosiga brevicollis (UniParc:*UPI000163B8DD); *Hydra magnipapillata (UniParc:*UPI0002B4BB35); *Capitella teleta* (*UniParc:*UPI00032C0488); *Aplysia californica* (*UniParc*:UPI0003593758); *Branchiostoma floridae* (*UniParc*:UPI0001861DC4); *Danio rerio* (*UniParc*:UPI000000BC41), *Xenopus tropicalis* (*UniParc*:UPI00004D064B); *Chrysemys picta bellii* (*UniParc*:UPI000388ED85); *Taeniopygia guttata* (*UniParc*: UPI000194BDA7); *Gallus gallus* (*UniParc*:UPI000044358E); *Melopsittacus undulates* (*UniParc*:UPI000383468D); *Amblyomma variegatum* (*UniParc*:UPI0002009EA1); *Caligus clemensi* (*UniParc*:UPI000198AD9A); *Lepeophtheirus salmonis* (*UniParc*:UPI000198A411); *Caligus rogercresseyi* (*UniParc*:UPI000198A05F) *Daphnia pulex* (*UniParc*:UPI0001FEB3EB); *Acyrthosiphon pisum* (*UniParc*:UPI0002061392); *Riptortus pedestris* (*UniParc:*UPI00032B4F83), *Pediculus humanus corporis* (*UniParc*:UPI000186E225); *Tribolium castaneum* (*UniParc*:UPI0000D56980); *Dendroctonus ponderosae* (*UniParc*:UPI00020FF929); *Bombyx mori* (*UniParc*:UPI0000E5D066); *Helicoverpa armigera* (*UniParc*:UPI00019203AC); *Danaus plexippus* (*UniParc*:UPI000239C638); *Nasonia vitripennis* (*UniProt*:UPI00015B411E); *Camponotus floridanus* (*UniProt*:UPI0001E7B62E); *Megachile rotundata* (*UniParc*:UPI000258DA7A); *Bombus terrestris* (*UniParc*:UPI00021A85D9); *Bombus impatiens* (*UniParc*:UPI00022CAAA0); *Apis mellifera (UniParc:UPI0000DB7C8B)*; *Aedes aegypti* (*UniParc*:UPI0000D76E88); *Culex quinquefasciatus* (*UniParc*:UPI00016D8A73); *Ceratitis capitata* (*UniParc*:UPI00032984C4); *Drosophila willistoni* (*UniParc*:UPI00017D8E71); *Drosophila grimshawi* (*UniParc*:UPI00017C6DDE); *Drosophila mojavensis* (*UniParc*:UPI00017C7884); *Drosophila virilis* (*UniParc*:UPI00017D4931); *Drosophila melanogaster* (*UniParc*:UPI000000741C); *Ovis aries* (*RefSeq*: XDA75500.1); *Mus musculus* (*RefSeq*: XP_006515229.1); *Oryctolagus cuniculus* (*RefSeq*: XP_051688519.2); *Pan troglodytes* (*RefSeq*: PNI66831.1); *Manis pentadactyla* (*RefSeq*: KAI5178856.1).

All sequences were aligned using MUSCLE (*80*) with default parameters. The resulting alignment was manually inspected, and conserved residues and motifs were visualized using ESPript 3.2(*81, 82*).

### Figures

Figures were prepared using ChimeraX 1.9 and Affinity Designer 2.

