## Supplementary material for "Structural basis for start codon selection by eIF5-mimic proteins"

**This PDF file includes:**

Figs. S1 to S12

Tables S1 to S3

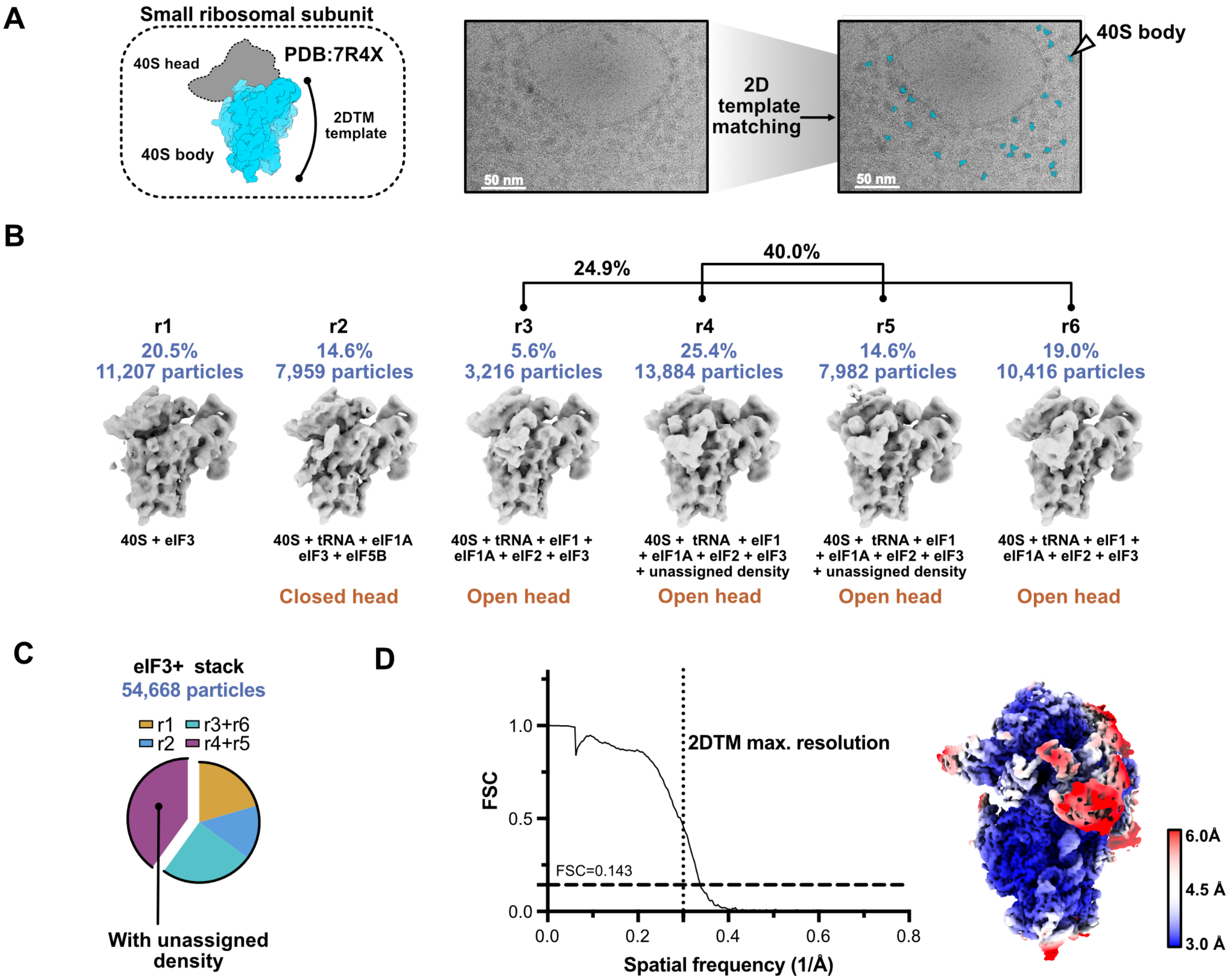


**Fig. S1. Identification of translation pre-initiation complexes in BSC-1 lysates.**
**(A)** Schematic representation of the template used for 2D template matching (2DTM), representative micrograph of BSC-1 cellular lysate and an illustration of 2DTM detections (blue). Template matching was carried out using the 40S body from PDB 7R4X. **(B)** Pre-initiation complex (PIC) classes obtained after 3D classification of 2DTM-identified particles (8× binned data). Prior particle sorting removed 80S and non-ribosomal particles. **(C)** Pre-initiation complexes account for ~11% of all particles detected in BSC-1 lysate. Among these, ~40% of the particles adopt an open-head conformation containing an additional unassigned density, which we identified as a member of the 5MP protein family. **(D)** Fourier shell correlation (FSC) curve showing the global resolution. The resolution limit imposed during 2DTM is indicated by the dotted line (~3.2 Å). A local resolution map of the 48S•5MP reconstruction is shown, with higher-resolution regions indicated in blue (~3.0 Å) and lower-resolution regions in red (~6.0 Å). The map was denoised using RELION.

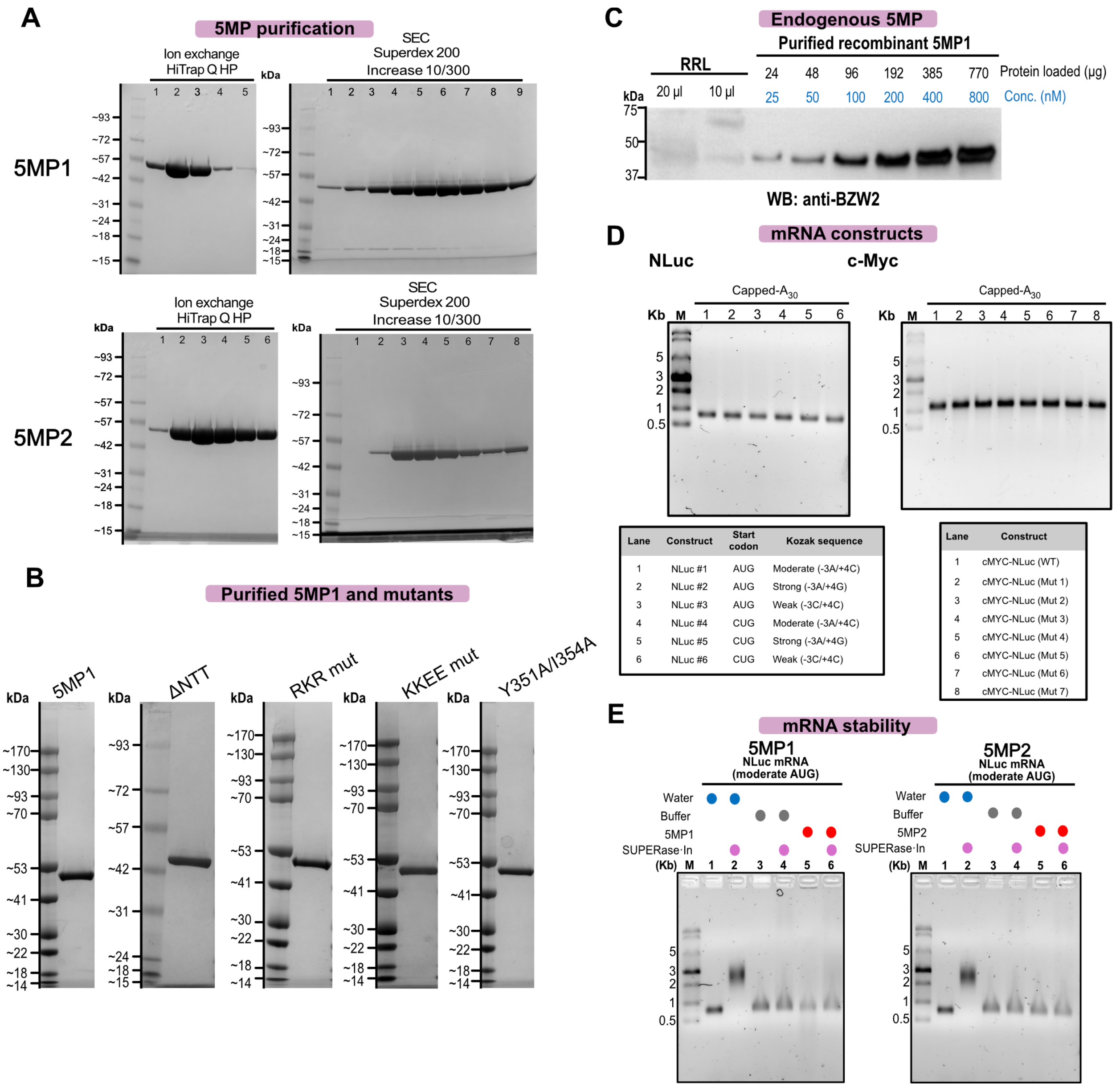


**Fig. S2. Preparation and validation of recombinant 5MP proteins and reporter mRNAs.** **(A)** Recombinant eIF5-mimic proteins 5MP1 (top) and 5MP2 (bottom) were purified by His-tag affinity chromatography, followed by ion-exchange separation on a HiTrap Q HP column, and a final purification step by size-exclusion chromatography using a Superdex 200 Increase 10/300 column. SDS-PAGE results are shown. For 5MP1, fractions 6-9 from the size-exclusion step were pooled, quantified, and used for translation assays and cryo-EM. For 5MP2, fractions 3-6 were collected for subsequent experiments. **(B)** SDS-PAGE of purified 5MP1 and mutants used for functional assays (2 µg per well): the NTT-deletion residues 1-25 mutant (ΔNTT), the RKR(17-19)-to-AAA mutant (RKR mut), the KKEE(309-312)-to-EEQQ mutant (KKEE mut), and the Y351A/I354A double mutant. **(C)** The concentration of endogenous 5MP in rabbit reticulocyte lysate (RRL) is estimated to be < 25 nM. Titrations were performed using recombinant 5MP1, and detection was performed with an anti-BZW2 antibody. **(D)** Capped mRNA constructs with $A_{30}$ tails were produced by *in vitro* transcription for translation assays and cryo-EM. **(E)** Purified 5MP1 and 5MP2 showed no detectable RNase contamination: mRNA stability in the presence of purified 5MPs was assessed by incubating the moderate-AUG reporter mRNA (the construct used for cryo-EM) for 1 hour at 37 °C, *i.e.*, substantially longer than the typical reaction time in this work (~20 minutes).

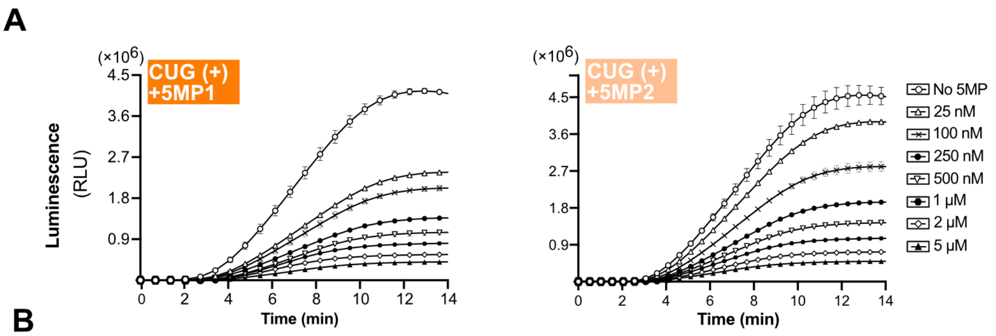


**B**

| 5MP1 | | | | | | |
|---|---|---|---|---|---|---|
| Start codon | AUG | AUG | AUG | CUG | CUG | CUG |
| Kozak context | Strong (−3A/+4G) | Moderate (−3A/+4C) | Weak (−3C/+4C) | Strong (−3A/+4G) | Moderate (−3A/+4C) | Weak (−3C/+4C) |
| $IC_{50}$(nM, mean ± SEM) | > 5 μ M | 701±66 | 114±21 | 35±4 | 11±2 | 11±1 |
| **5MP2** | | | | | | |
| Start codon | AUG | AUG | AUG | CUG | CUG | CUG |
| Kozak context | Strong (−3A/+4G) | Moderate (−3A/+4C) | Weak (−3C/+4C) | Strong (−3A/+4G) | Moderate (−3A/+4C) | Weak (−3C/+4C) |
| $IC_{50}$(nM, mean ± SEM) | > 5 μ M | 764±87 | 410±31 | 131±10 | 79±3 | 82±6 |

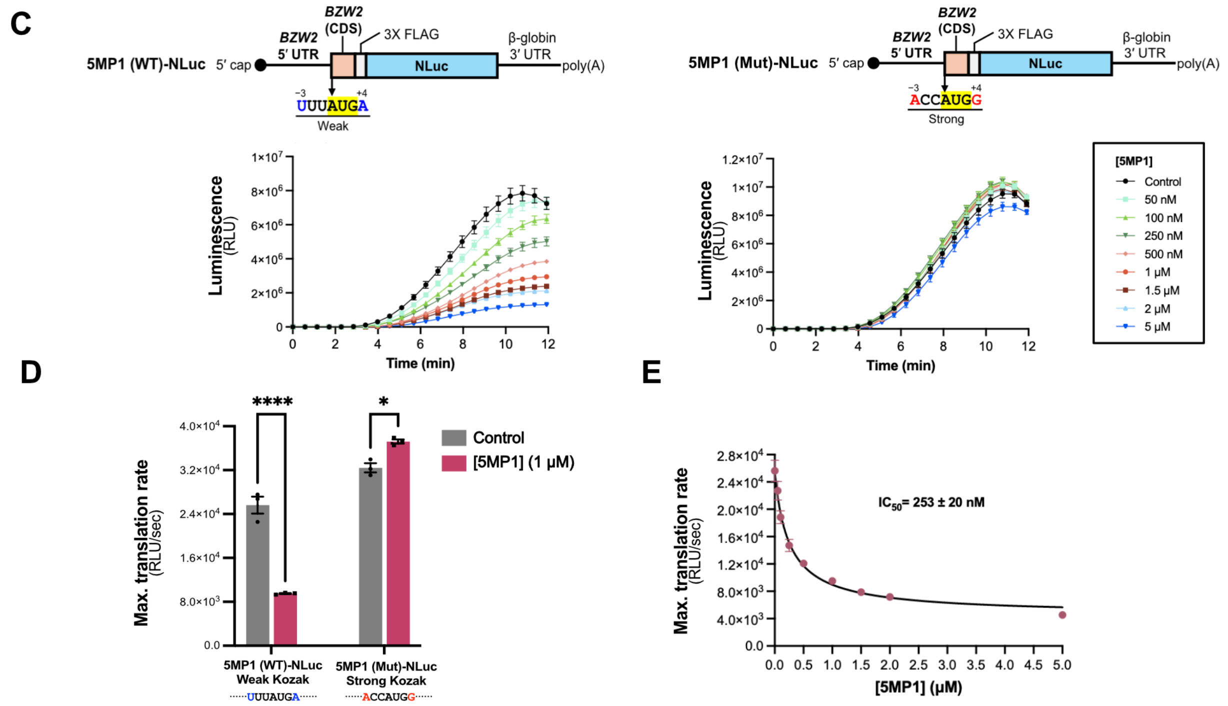


**Fig. S3. 5MP-Mediated Translation Inhibition and Self-Regulation.**

**(A)** Time progress curves for strong Kozak CUG initiation at increasing 5MP concentrations (25 nM, 100 nM, 250 nM, 500 nM, 1μM, 2 μM and 5 μM) **(B)** $IC_{50}$ values for translation inhibition of NLuc constructs with different start codons by 5MP1 and 5MP2 in rabbit reticulocyte lysate (RRL); see additional data in main-text Fig. 1. **(C)** Schematic representation of the mRNA construct containing the 5′UTR and coding sequence (CDS) of the 5MP1 (*BZW2*) transcript, followed by a 3×FLAG tag and a nanoluciferase (NLuc) reporter. To assess self-regulation, constructs were generated with either the native translation initiation site (wild type; WT), which contains an AUG codon in a weak Kozak context (−3U/+4A, left), or a mutant version (Mut) in which AUG is placed in a strong Kozak context (−3A/+4G, right). Time-course analysis of *in vitro* RRL translation with increasing concentrations of 5MP1 for the native (WT; left) and mutant (Mut; right) constructs, demonstrating the self-regulatory effect of 5MP1. **(D)** Bar chart showing the effect of 5MP1 (1 μM) on translation of the native (WT; weak Kozak, left) and mutant (Mut; optimal Kozak, right) 5MP1-NLuc mRNAs. **(E)** Translation inhibition curve showing the 5MP1 inhibitory concentration ($IC_{50}$) for the mRNA carrying the WT-5MP1 start codon in RRL. Data represent mean ± SEM (n=3) (C) and mean ± SD (n=3) (D, E). Statistical significance on maximum translation rates was assessed by two-way ANOVA; ****P < 0.0001, ***P < 0.0002, **P < 0.0021, ns = not significant.

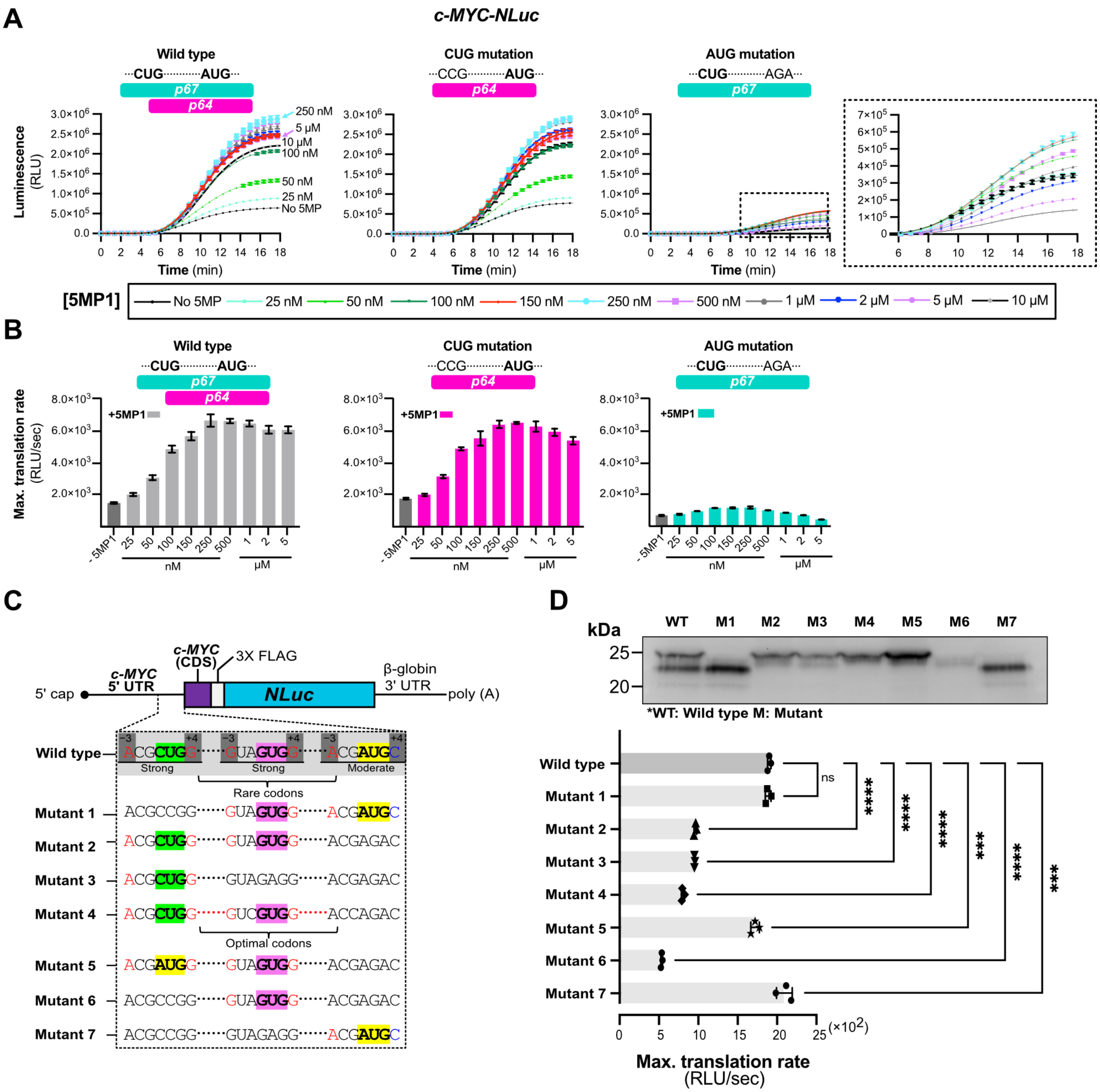


**Fig. S4. 5MP1 regulation of translation from *the c-Myc* 5′ UTR and inefficient upstream CUG initiation**

**(A)** Time course analysis of in vitro RRL translation of the c-Myc-NLuc wild type construct, the CUG-mutant (Mut 1), and the AUG-mutant ( Mut 2). (A, right) Close-up view of the translation curves. **(B)** Schematic representation of the c-Myc reporter constructs. Previously reported translation initiation sites included an upstream CUG within a strong Kozak context and a downstream AUG within a moderate Kozak context. Additional constructs were designed to modify initiation efficiency by mutating the CUG codon, replacing rare codons between the CUG and AUG start codons with frequent codons, or mutating the GUG codon between CUG and AUG. **(C)** Schematic of the *c-Myc* 5′ UTR reporter showing the in-frame initiation sites, Kozak contexts, and mutants. **(D)** Translation products detected by anti-FLAG western blotting (top) and maximum translation rates for *c-Myc* wild-type 5′ UTR and the mutants (bottom). Data represent mean ± SEM (n=3) (A, B) and mean ± SD (n=3) (D). Statistical significance of maximum translation rates was assessed by one-way ANOVA; ****$P < 0.0001$, ***$P < 0.0002$, **$P < 0.0021$; ns, not significant.

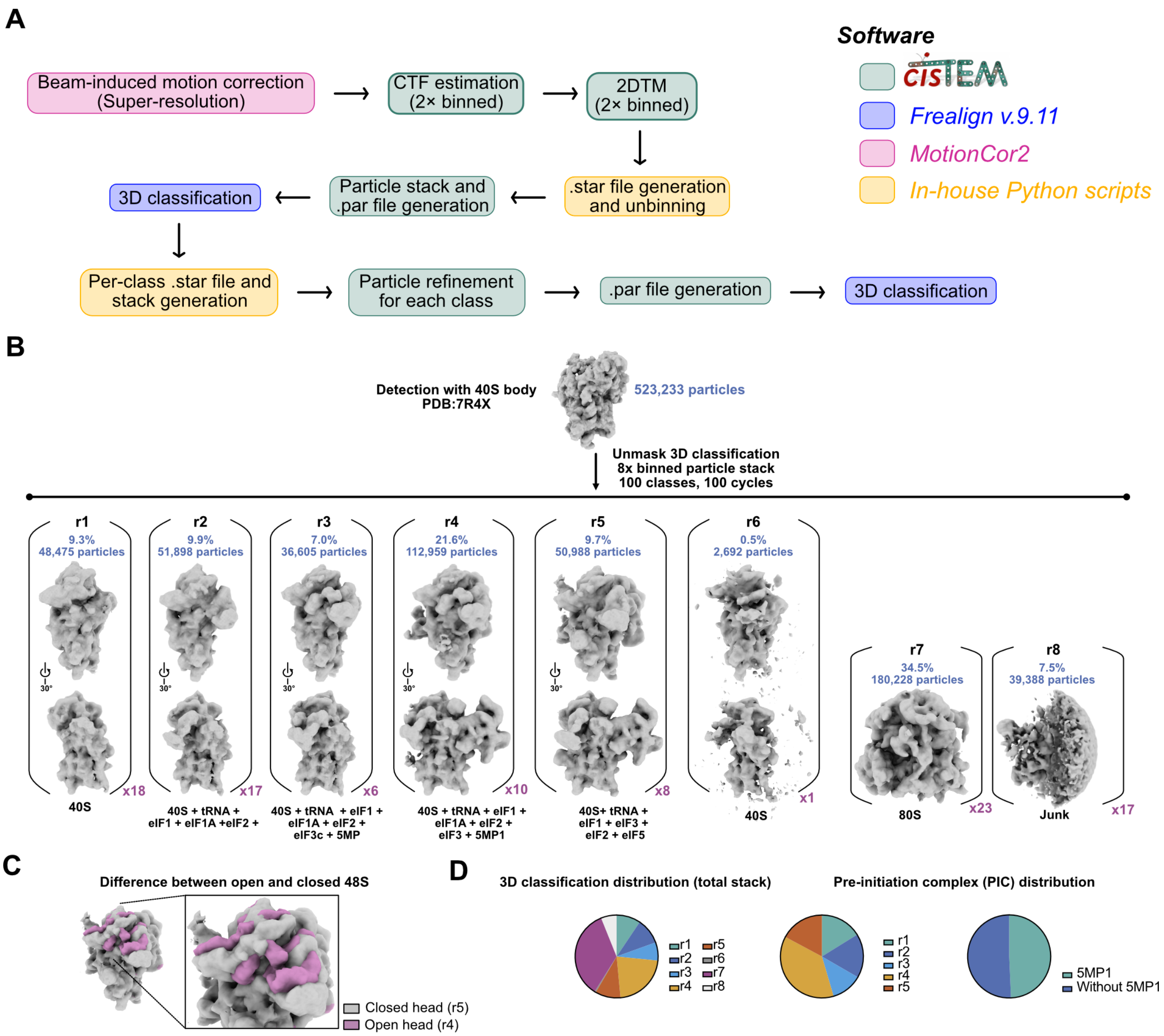


**Fig. S5. Cryo-EM image processing and classification workflow for RRL supplemented with recombinant 5MP1.**

**(A)** Beam-induced motion correction was performed on dose-fractionated movies using MotionCor2 at super-resolution. Contrast transfer function (CTF) parameters were estimated from 2× binned micrographs using *cis*TEM, followed by 2D template matching (2DTM) on binned data. Particle coordinates were unbinned using in-house Python scripts, and corresponding particle stacks and .par files were generated. Unbinning refers to generating particle stacks from the original micrographs at the original pixel size. Initial 3D classification was carried out in Frealign v9.11. For each resulting class, per-class particle stacks and .star files were generated and subjected to independent particle refinement. Refined parameters were then used to create updated .par files for subsequent rounds of focused or global 3D classification (see Methods). Software packages used at each step are indicated by color coding. In-house scripts are available in YAFW: https://github.com/GrigorieffLab/yafw. **(B)** 3D maximum-likelihood classification was performed in Frealign v9.11. Subclassification was carried out on binned data. **(C)** Structural differences between the open- and closed-head conformations are shown for classes r4 and r5, respectively. **(D)** Distribution of particle classes across the whole dataset and pre-initiation complexes (PICs), which correspond to ~58% of particles. Within the PIC population, ~50% of particles contain 5MP1. Among these, 25% correspond to 43S PICs with only the eIF3c subunit visible (class r3), and the rest correspond to 48S complexes containing 5MP1 (class r4). A closed-head 48S conformation (class r5) accounts for 9.7% of particles.

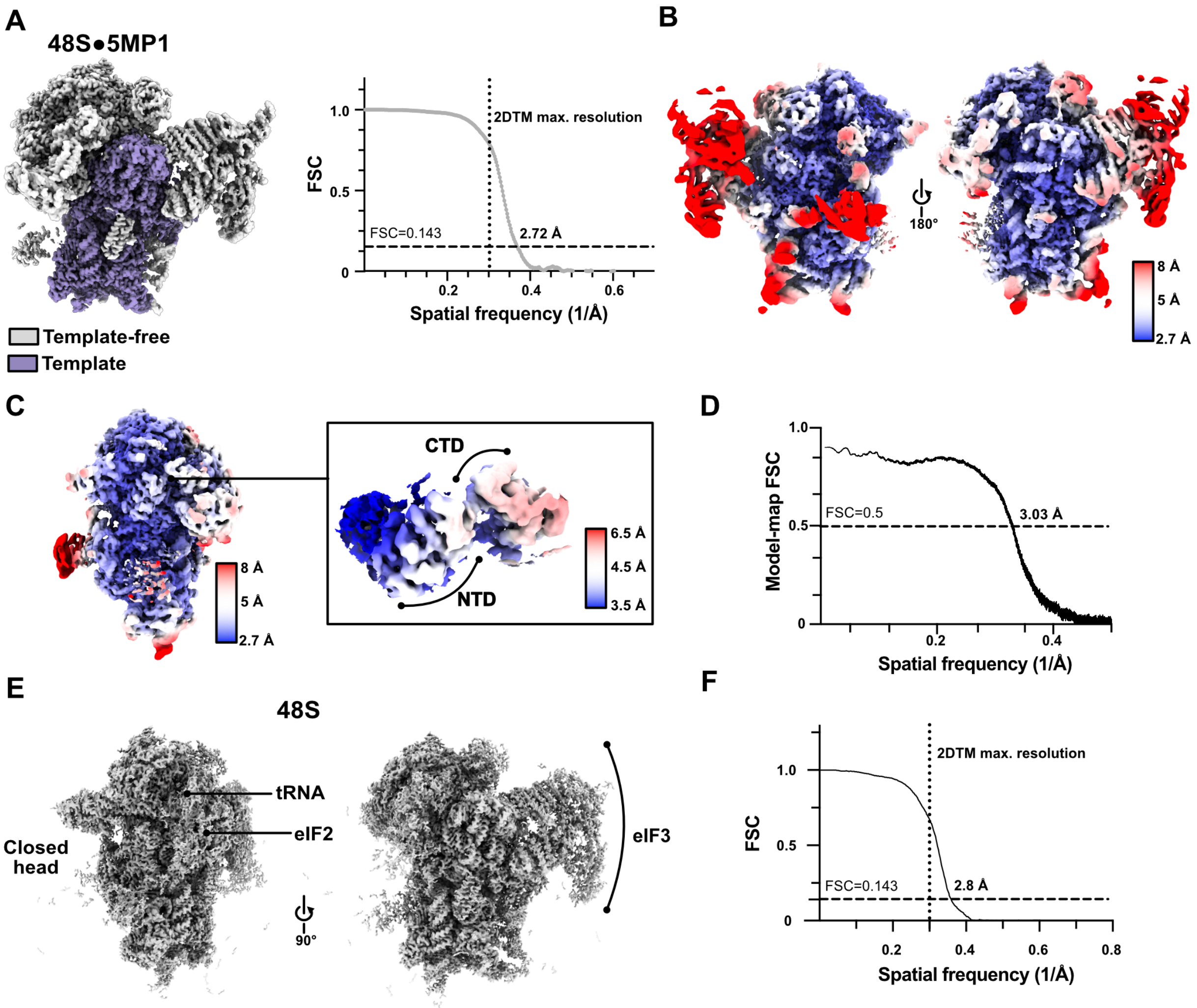


**Fig. S6. Assessment of average resolution, template bias and local resolution of the 48S•5MP1 and 48S pre-initiation complex.**

**(A)** Reconstruction of the 48S•5MP1 initiation complex (class r4) following manual refinement and post-processing with DeepEMenhancer. Regions included in the template are shown in purple, and omitted areas are in gray. The Fourier shell correlation (FSC) curve used for global resolution estimation is shown. Template matching was performed on 2× binned data (1.66 Å/pixel), imposing a resolution limit of 3.32 Å (right). Therefore, density features at resolutions higher than ~3.3 Å cannot be attributed to template bias. **(B-C)** Local resolution estimation of the 48S•5MP1 map and close-up view of the local resolution for 5MP1; the N-terminal domain (NTD) and C-terminal domain (CTD) are labelled. **(D)** FSC between the model and the map was calculated for the model refined against the map. **(E)** Cryo-EM density map of the closed-head 48S complex lacking 5MP1, refined to 2.8 Å resolution. The reconstruction shows density for the initiator $tRNA^{Met}$, the eIF2 ternary complex, and eIF3. Density corresponding to mRNA in the mRNA tunnel is also present (see Fig. 4 and Extended Data Figure 11). Two orthogonal views are shown, highlighting the closed conformation of the 40S head relative to the body and the presence of eIF3. **(F)** Fourier shell correlation (FSC) curve for the global reconstruction of the closed-head 48S complex. The 2DTM resolution limit is indicated with a dotted line.

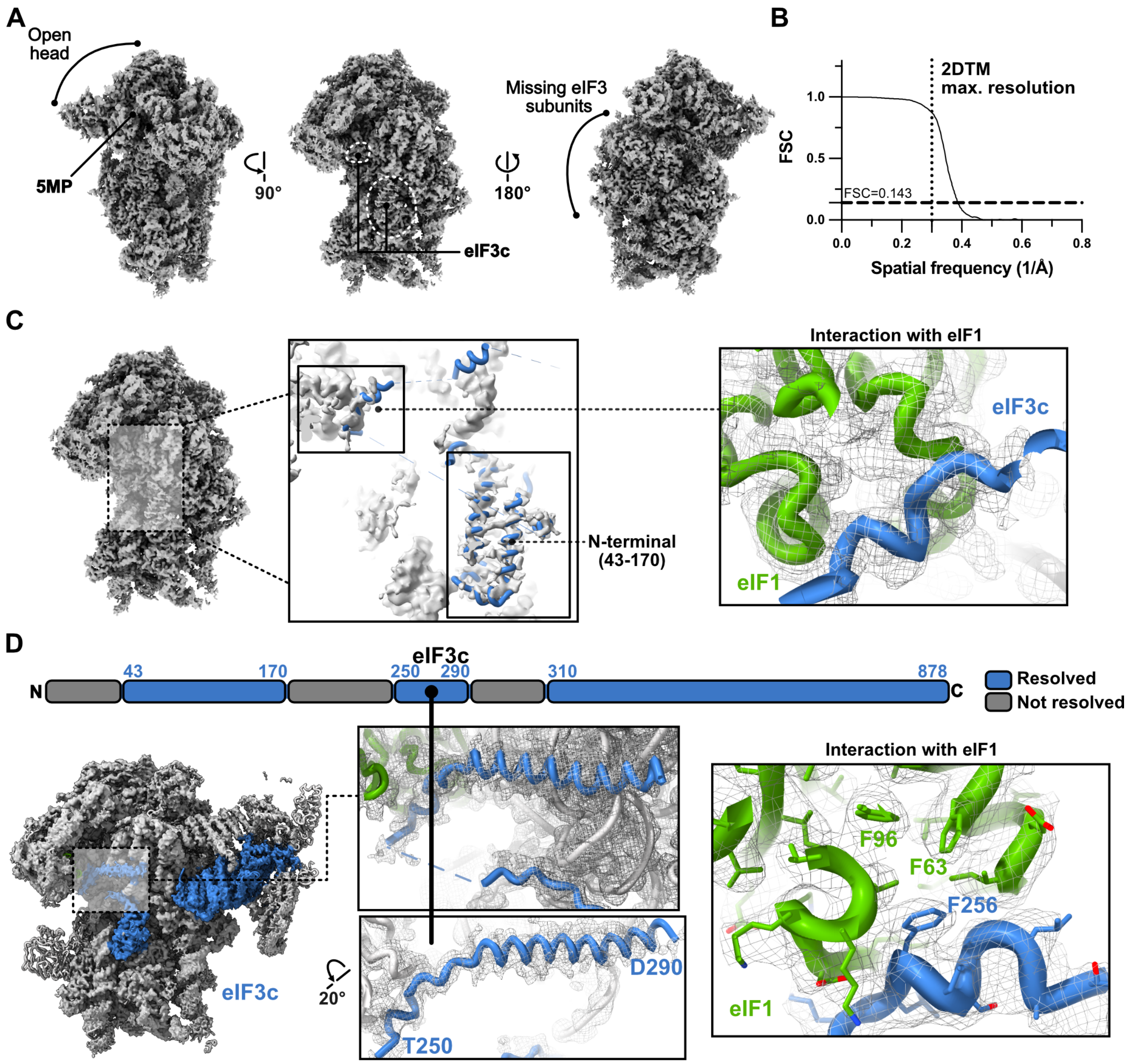


**Fig. S7. The N-terminal domain of eIF3c contacts eIF1.**

**(A)** 2.8 Å map of 5MP1 bound to a 48S-like pre-initiation complex showing densities consistent with the eIF3c N-terminal region. Density is observed for residues 43-260 of eIF3c, whereas no density is detected for the remaining eIF3 subunits. **(B)** Fourier shell correlation (FSC) curve for the reconstruction shown in panel A. The 2DTM resolution limit is indicated by a dotted line. **(C)** Densities corresponding to the eIF3c helix interacting with eIF1 (also see box panel D for reference). **(D)** A scheme and densities in the 48S complex with eIF3, showing the resolved segments of eIF3c in blue and eIF1 model in green. The interaction supports a role of the eIF3c N-terminal domain in 5MP1-mediated regulation of initiation stringency *(50)*. Residues T250–D290 (middle) were initially modeled using AlphaFold2 DB and refined using Isolde and Phenix. The structure revealed that the Phe256 of the FLKK motif 3 of eIF3c *(50)* is inserted into a hydrophobic pocket on the surface of eIF1 (right).

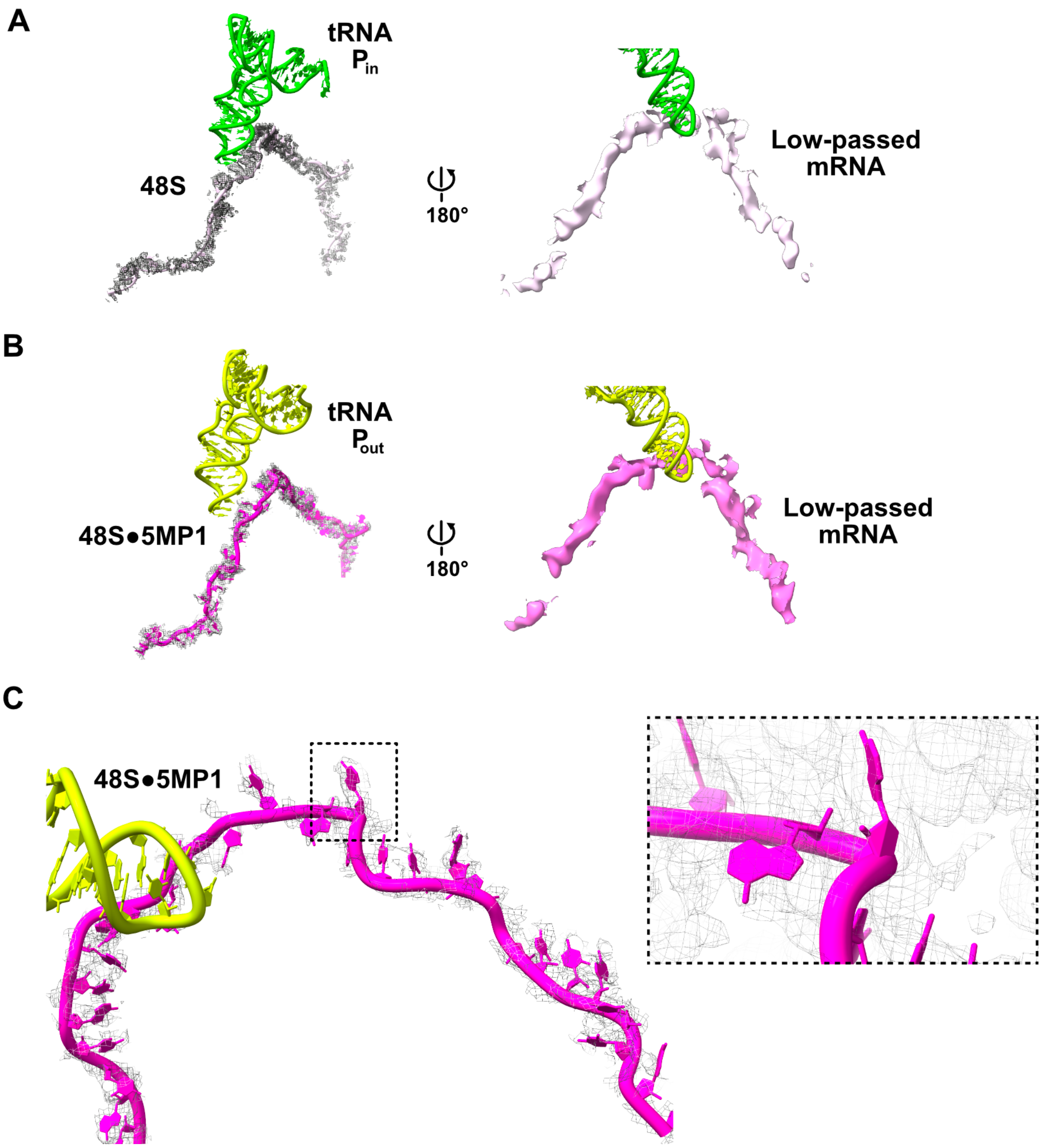


**Fig. S8. Scanning 48S•5MP1 initiation complexes exhibits continuous density for the mRNA.**

**(A)** In the 48S complex, mRNA density is continuous throughout the mRNA tunnel, and the initiator $tRNA^{Met}$ adopts a $P_{in}$ conformation, consistent with an arrested scanning state. **(B)** In the 48S•5MP1 complex, mRNA density is present within the mRNA tunnel, while the initiator $tRNA^{Met}$ adopts a $P_{out}$ conformation, consistent with a scanning state. **(C)** Close-up view of the mRNA tunnel in the 48S•5MP1 complex. Nucleotide densities are not well-resolved, consistent with increased heterogeneity in the scanning state, although density is present for some nucleotide positions, as shown in the close-up view, similar to those reported for the scanning 48S complex (EMD-17696; PDB 8PJ1)

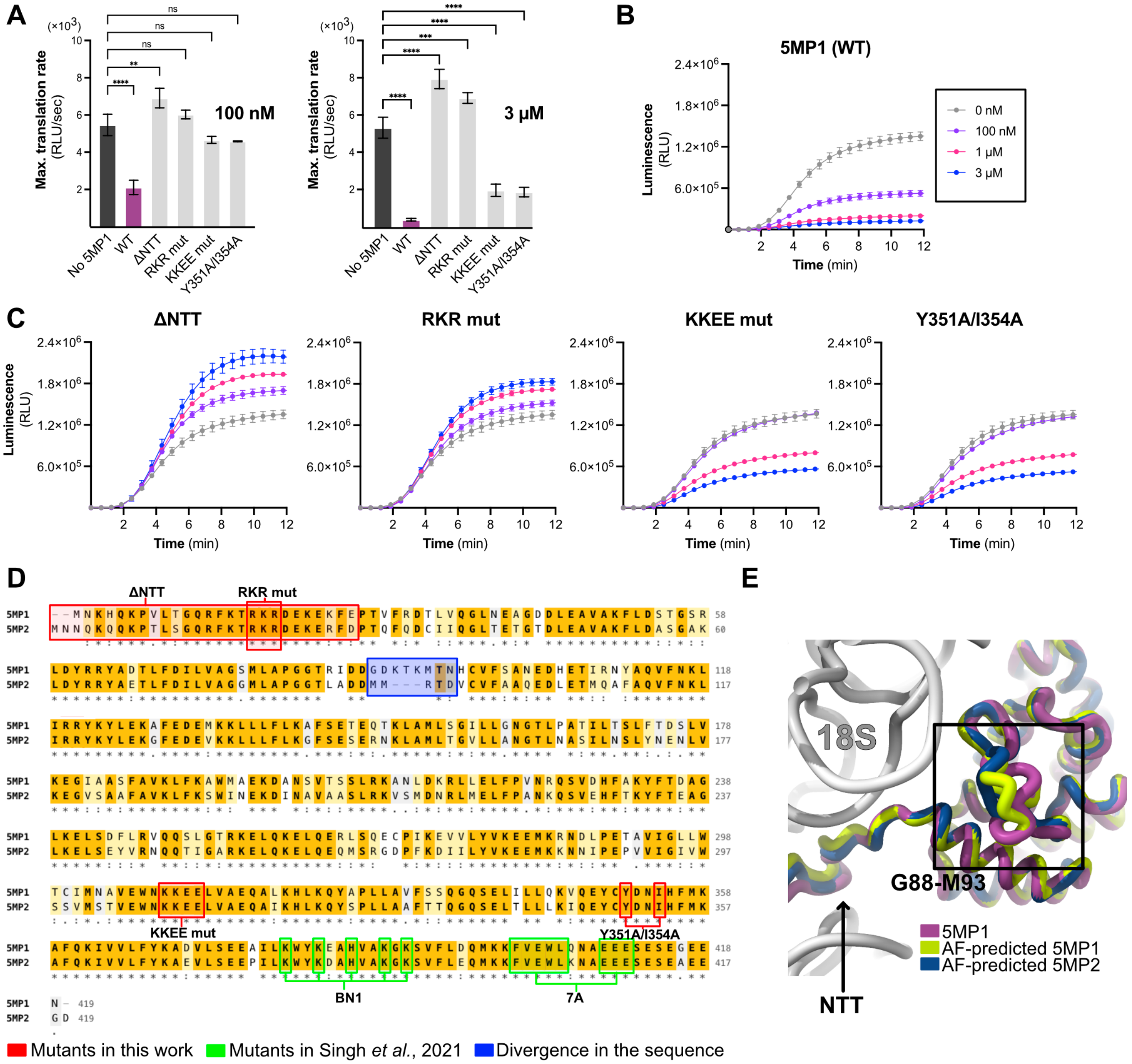


**Fig. S9. Effects of structure-guided mutations of 5MP1 on translation.**
**(A)** Maximal rates of translation of the transcript containing a CUG start codon in a strong Kozak context (see construct in Fig 1B), at 100 nM and 3 μM of 5MP1 WT or mutant proteins (mean ± SD, n=3). The rates are calculated from data in panels B and C. **(B)** Time progress curves for wild-type 5MP1 and **(C)** 5MP1 mutants (from left to right): the NTT-deletion residues 1-25 mutant (ΔNTT), the RKR(17-19)-to-AAA mutant (RKR mut), the KKEE(309-312)-to-EEQQ mutant ( KKEE mut), and the Y351A/I354A double mutant, at 100 nM, 1 μM, and 3 μM (mean ± SEM, n=3). Statistical significance was assessed by one-way ANOVA; ****P < 0.0001, ***P < 0.0002, **P < 0.0021, ns = not significant. **(D)** Sequence alignment of 5MP1/BZW2 (UniProt: Q9Y6E2) and 5MP2/BZW1 (UniProt: Q7L1Q6) performed using Clustal Omega. Mutations generated in this study are boxed in red, mutations from Singh et al., 2021 are boxed in green, and the most divergent sequence region is boxed in blue (Gly88–Asn95). **(E)** Close-up view of the divergent sequence mapped onto the structure (with the divergent Gly88-Met93 region clearly visible). This

region is located in the N-terminal domain (NTD). Our 5MP1 model is shown in purple, the AlphaFold prediction for 5MP1 (Q9Y6E2) in yellow-green, and for 5MP2 (Q7L1Q6) in navy blue.

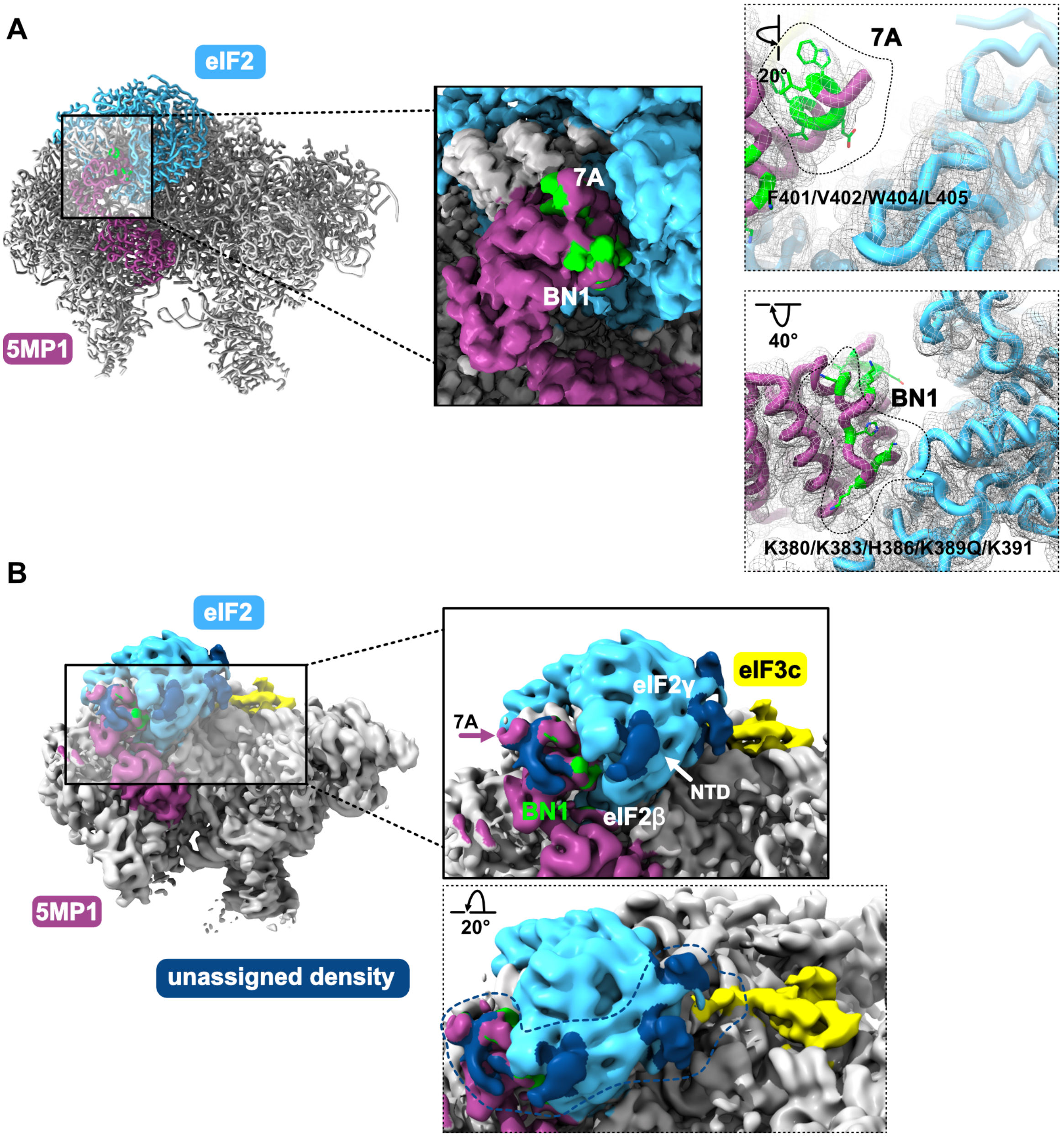


**Fig. S10. Identification of 7A and BN1 mutations in 5MPs and of unassigned densities that potentially interact with the amino acids altered in these mutants.**

**(A)** Identification of the mutations by Asano's group (Singh et al., 2021), BN1 and 7A, mapped onto our model and cryo-EM density map. Both mutation clusters (green) are located at the interface with eIF2β **(B).** Unassigned densities (dark blue) in an 8 Å low-passed filtered map displayed at a low contour level. These densities could be attributed to eIF2β N-terminal tail (densities near 5MP1 and eIF2β) or to eIF3c N-terminal (all densities shown in dark blue). Noise can not be excluded as an explanation for these densities. 5MP1 is shown in purple, eIF2 in light blue and eIF3c in yellow. 7A and BN1 mutations are shown in green. NTD stand for N-terminal domain, here identified for eIF2β.

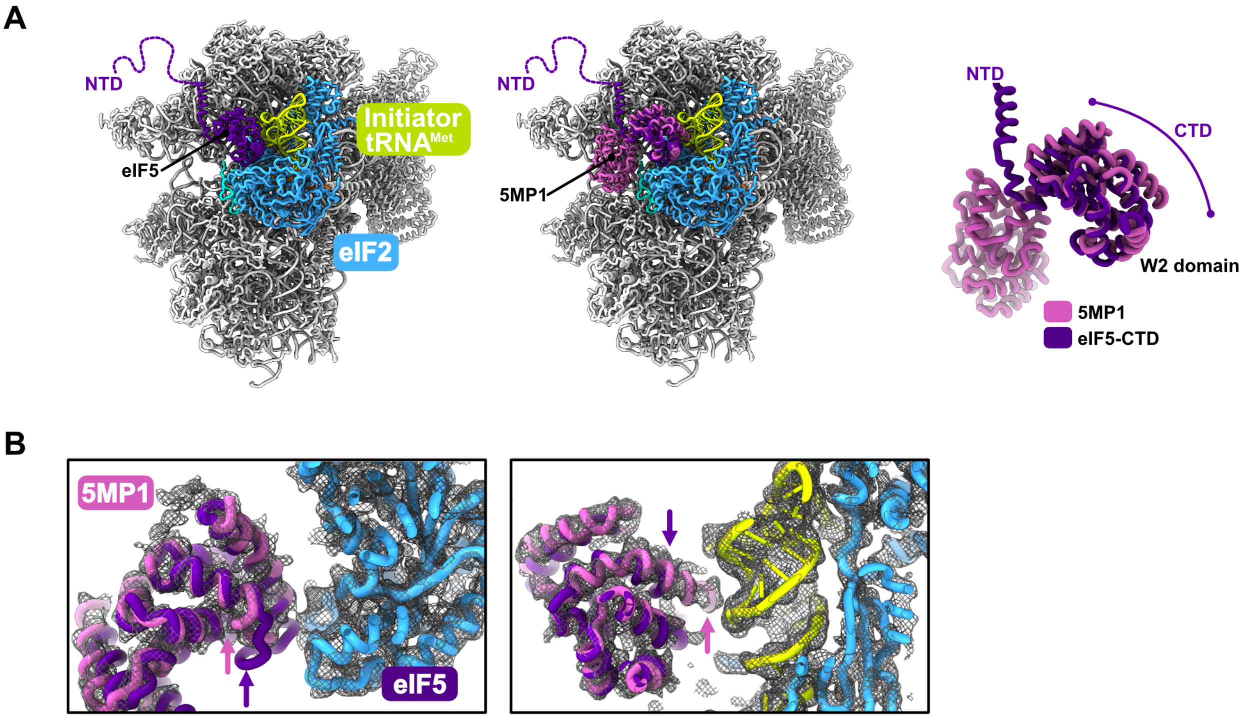


**Fig. S11. Comparison of the CTDs from 5MP1 and from a predicted model of eIF5.**
**(A)** Superposition of the AlphaFold2-predicted structure of the eIF5 C-terminal domain (CTD; deep purple) onto the 48S•5MP1 cryo-EM structure (this work), aligned on 5MP1 CTD. Modeled densities are shown for 5MP1 in purple, eIF2 in blue, eIF1 in orange, eIF1A in turquoise, and initiator $tRNA^{Met}$ in green. The overlay suggests that the eIF5 CTD (left panel) could occupy a similar position within the 48S as the 5MP1 CTD (middle and right panels), consistent with potential competition between eIF5 and 5MP1 for binding at this site. **(B)** Close-up view of the eIF5 CTD and 5MP1 models, along with the 48S•5MP1 cryo-EM densities for eIF2β and initiator $tRNA^{Met}$, suggesting distinct molecular contacts.

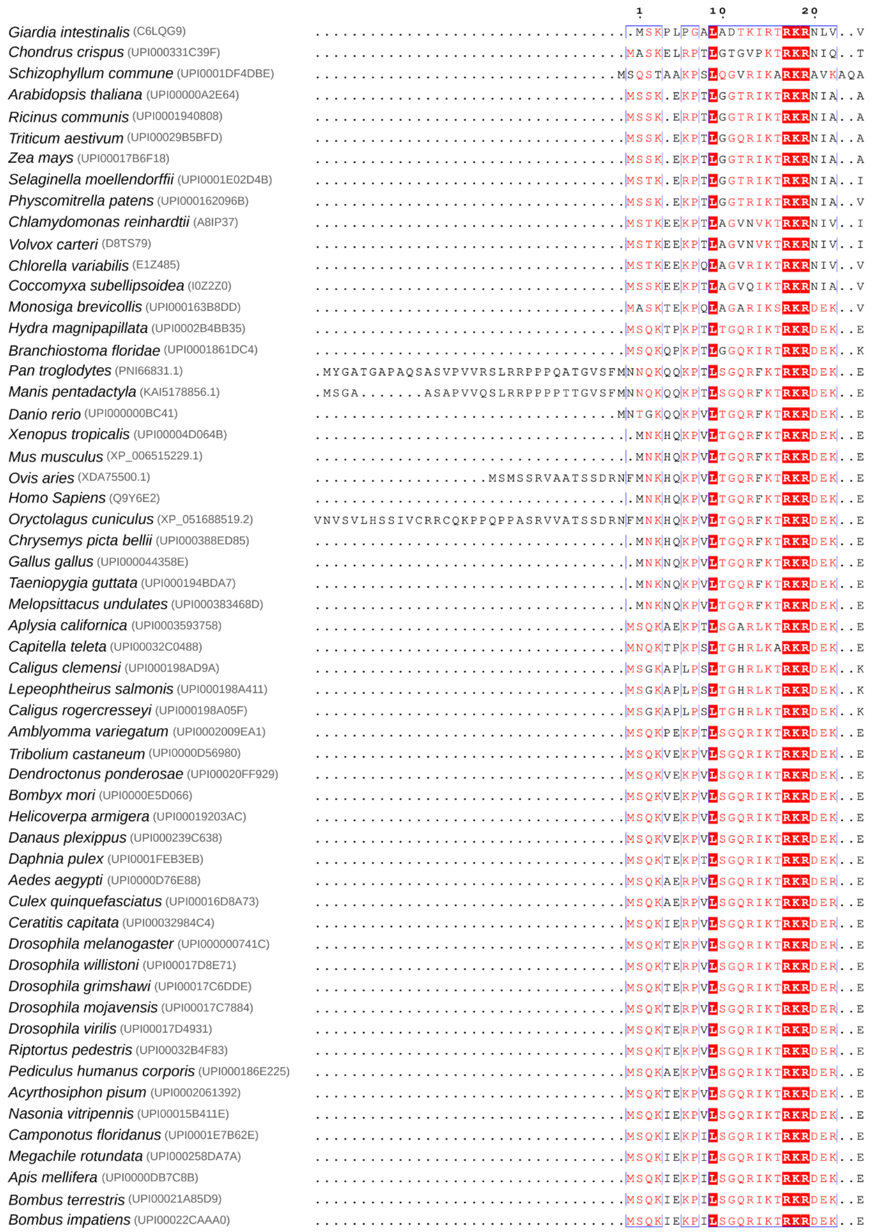


**Fig. S12. Conservation of the 5MP1 N-terminal tail and RKR motif across eukaryotes.** Multiple sequence alignment of the 5MP1 N-terminal region (residues 1–20; human 5MP1 numbering, UniProt Q9Y6E2) across 57 species, ranging from unicellular eukaryotes to vertebrates, generated with MUSCLE and rendered with ESPript 3.2. Species names are shown with the corresponding UniProt/UniParc/RefSeq accession used for the alignment. Identical residues are shaded red. The Arg17-Lys18-Arg19 (RKR) motif, which contacts the mRNA within the 40S mRNA tunnel (Fig. 5E), is among the most highly conserved elements.

**Table S1. Near-Cognate Start Codons in the *c-MYC* 5′UTR (nt 1–408)**

| # | Position | Codon | Frame | Codons upstream | Kozak | | | Sequence |
|---|---|---|---|---|---|---|---|---|
| | | | | | -3 | +4 | Strength | |
| 1 | 7 | CUG | In-frame | 134 | U | U | weak | ACUCG[CUG]UAGUA |
| 2 | 81 | CUG | Out-of-frame | 109 | G | C | moderate | CAGAG[CUG]CGCUG |
| 3 | 86 | CUG | Out-of-frame | 107 | G | C | moderate | CUGCG[CUG]CGGGC |
| 4 | 97 | CUG | In-frame | 104 | G | G | strong | GCGUC[CUG]GGAAG |
| 5 | 136 | CUG | In-frame | 91 | C | G | moderate | CGCCU[CUG]GCCCA |
| 6 | 153 | CUG | Out-of-frame | 85 | C | A | weak | UCCCG[CUG]AUCCC |
| 7 | 231 | CUG | Out-of-frame | 59 | G | G | strong | UUGCA[CUG]GAACU |
| 8 | 287 | CUG | Out-of-frame | 40 | A | C | moderate | CUAUU[CUG]CCCAU |
| 9 | 315 | CUG | Out-of-frame | 31 | C | C | weak | CGCCG[CUG]CCAGG |
| 10 | 333 | CUG | Out-of-frame | 25 | U | A | weak | CUUCU[CUG]AAAGG |
| 11 | 353 | CUG | Out-of-frame | 18 | C | C | weak | UGCAG[CUG]CUUAG |
| 12 | 364 | CUG | In-frame | 15 | A | G | strong | AGACG[CUG]GAUUU |
| 13 | 182 | UUG | Out-of-frame | 75 | C | C | weak | AACCC[UUG]CCGCA |
| 14 | 201 | UUG | Out-of-frame | 69 | A | C | moderate | AAACU[UUG]CCCAU |
| 15 | 226 | UUG | In-frame | 61 | A | C | moderate | GCACU[UUG]CACUG |
| 16 | 295 | UUG | In-frame | 38 | C | G | moderate | CCCAU[UUG]GGGAC |
| 17 | 347 | UUG | Out-of-frame | 20 | U | C | weak | UCUCC[UUG]CAGCU |
| 18 | 58 | GUG | In-frame | 117 | A | G | strong | CUAGG[GUG]GAAGA |
| 19 | 382 | GUG | In-frame | 9 | G | G | strong | GGGUA[GUG]GAAAA |

**Table S2. Cryo-EM data collection, 2DTM results, and model refinement and validation statistics.**

| **Data Collection** | **BSC-1** | **RRL + mRNA + 5MP1** |
|---|---|---|
| Microscope | Titan Krios | Titan Krios |
| Voltage (kV) | 300 | 300 |
| Magnification | 105000 | 105000 |
| Pixel size (Å) | 0.83 | 0.83 |
| Detector | K3 | K3 |
| Defocus range (μm) | 1.0-1.5 | 1.0-1.5 |
| Total electron exposure ($e^{-}Å^{-2}s^{-}$) | ~30 | ~30 |
| Data collection software | SerialEM | SerialEM |
| Micrograph collected (no.) | 162 583 | 104 673 |
| EMPIAR ID | 13853/13577 | 13852 |
| **Sample Geometry** | | |
| Average sample thickness (nm)* | 143 ± 13 | 148 ±12 |
| **2DTM** | | |
| Total number of particles (no.) | 1 573 843 | 523 233 |
| Average SNR* | 8.1 | 8.0 |
| **Model refinement** | | **48S•5MP1** |
| EM Data Bank and PDB identifiers | | EMD-78242, 37KM |
| Number of particles (no.) | | 112959 |
| Average map resolution (Å) (FSC threshold=0.143) | | 2.7 |
| Map resolution range (Å) | | 2.7-16 |
| Symmetry imposed | | *C1* |
| **Atomic model refinement** | | |
| Initial model used (PDB code) | | Methods |
| Model resolution (Å) | | 2.8 |
| Map sharpening B-factor ($Å^2$) | | -50 |
| CCmask | | 0.84 |
| Model composition, no. atoms | | 112238 |
| RNA residues | | 1813 |
| Protein residues | | 9536 |
| **B-factors ($Å^2$)** | | |
| **RNA residues** | | 38.44/398.50/113.12 |
| Proteins | | 37.30/487.39/159.19 |
| **Root-mean square deviations** | | |
| Bond lengths (Å) | | 0.015 |
| Bond angles (°) | | 1.126 |
| **Validation** | | |
| MolProbity score | | 2.20 |
| Clashscore, all atoms | | 6.81 |
| Good sugar pucker (%) | | 98.62 |
| **Ramachandran plot** | | |
| Favored (%) | | 94.0 |

*Over 10 representative micrographs

**Table S3: DNA sequences of NLuc reporters and constructs with *c-Myc* 5′ UTR.**

| Construct | Sequence (5′-3′) |
|---|---|
| **Strong AUG (−3A/+4G)** | acacttgcttttgacacaactgtgtttacttgcaatcccccaaaacagacaatggatcatcatcatcatcatggctcgagcggcgtcttcacactcgaagatttcgttg<br>gggactggcgacagacagccggctacaacctggaccaagtccttgaacagggaggtgtgtccagtttgtttcagaatctcggggtgtccgtaactccgatccaa<br>aggattgtcctgagcggtgaaaatgggctgaagatcgacatccatgtcatcatcccgtatgaaggtctgagcggcgaccaaatgggccagatcgaaaaaattttt<br>aaggtggtgtaccctgtggatgatcatcactttaaggtgatcctgcactatggcacactggtaatcgacggggttacgccgaacatgatcgactatttcggacggc<br>cgtatgaaggcatcgccgtgttcgacggcaaaaagatcactgtaacagggaccctgtggaacggcaacaaaattatcgacgagcgcctgatcaaccccgacg<br>gctccctgctgttccgagtaaccatcaacggagtgaccggctggcggctgtgcgaacgcattctggcgtaagatctttttccctctgccaaaaattatggggacat<br>catgaagccccttgagcatctgacttctggctaataaaggaaatttattttcattgc |
| **Moderate AUG (−3A/+4C)** | acacttgcttttgacacaactgtgtttacttgcaatcccccaaaacagacaatgcatcatcatcatcatcatggctcgagcggcgtcttcacactcgaagatttcgttg<br>gggactggcgacagacagccggctacaacctggaccaagtccttgaacagggaggtgtgtccagtttgtttcagaatctcggggtgtccgtaactccgatccaa<br>aggattgtcctgagcggtgaaaatgggctgaagatcgacatccatgtcatcatcccgtatgaaggtctgagcggcgaccaaatgggccagatcgaaaaaattttt<br>aaggtggtgtaccctgtggatgatcatcactttaaggtgatcctgcactatggcacactggtaatcgacggggttacgccgaacatgatcgactatttcggacggc<br>cgtatgaaggcatcgccgtgttcgacggcaaaaagatcactgtaacagggaccctgtggaacggcaacaaaattatcgacgagcgcctgatcaaccccgacg<br>gctccctgctgttccgagtaaccatcaacggagtgaccggctggcggctgtgcgaacgcattctggcgtaagatctttttccctctgccaaaaattatggggacat<br>catgaagccccttgagcatctgacttctggctaataaaggaaatttattttcattgc |
| **Weak AUG (−3C/+4C)** | acacttgcttttgacacaactgtgtttacttgcaatcccccaaaacagccaatgcatcatcatcatcatcatggctcgagcggcgtcttcacactcgaagatttcgttg<br>gggactggcgacagacagccggctacaacctggaccaagtccttgaacagggaggtgtgtccagtttgtttcagaatctcggggtgtccgtaactccgatccaa<br>aggattgtcctgagcggtgaaaatgggctgaagatcgacatccatgtcatcatcccgtatgaaggtctgagcggcgaccaaatgggccagatcgaaaaaattttt<br>aaggtggtgtaccctgtggatgatcatcactttaaggtgatcctgcactatggcacactggtaatcgacggggttacgccgaacatgatcgactatttcggacggc<br>cgtatgaaggcatcgccgtgttcgacggcaaaaagatcactgtaacagggaccctgtggaacggcaacaaaattatcgacgagcgcctgatcaaccccgacg<br>gctccctgctgttccgagtaaccatcaacggagtgaccggctggcggctgtgcgaacgcattctggcgtaagatctttttccctctgccaaaaattatggggacat<br>catgaagccccttgagcatctgacttctggctaataaaggaaatttattttcattgc |
| **Strong CUG (−3A/+4G)** | acacttgcttttgacacaactgtgtttacttgcaatcccccaaaacagacactggatcatcatcatcatcatggctcgagcggcgtcttcacactcgaagatttcgttg<br>gggactggcgacagacagccggctacaacctggaccaagtccttgaacagggaggtgtgtccagtttgtttcagaatctcggggtgtccgtaactccgatccaa<br>aggattgtcctgagcggtgaaaatgggctgaagatcgacatccatgtcatcatcccgtatgaaggtctgagcggcgaccaaatgggccagatcgaaaaaattttt<br>aaggtggtgtaccctgtggatgatcatcactttaaggtgatcctgcactatggcacactggtaatcgacggggttacgccgaacatgatcgactatttcggacggc<br>cgtatgaaggcatcgccgtgttcgacggcaaaaagatcactgtaacagggaccctgtggaacggcaacaaaattatcgacgagcgcctgatcaaccccgacg<br>gctccctgctgttccgagtaaccatcaacggagtgaccggctggcggctgtgcgaacgcattctggcgtaagatctttttccctctgccaaaaattatggggacat<br>catgaagccccttgagcatctgacttctggctaataaaggaaatttattttcattgc |
| **Moderate CUG** | acacttgcttttgacacaactgtgtttacttgcaatcccccaaaacagacactgcatcatcatcatcatcatggctcgagcggcgtcttcacactcgaagatttcgttg<br>gggactggcgacagacagccggctacaacctggaccaagtccttgaacagggaggtgtgtccagtttgtttcagaatctcggggtgtccgtaactccgatccaa |

| | |
|---|---|
| **(−3A/+4C)** | aggattgtcctgagcggtgaaaatgggctgaagatcgacatccatgtcatcatcccgtatgaaggtctgagcggcgaccaaatgggccagatcgaaaaaattttt<br>aaggtggtgtaccctgtggatgatcatcactttaaggtgatcctgcactatggcacactggtaatcgacggggttacgccgaacatgatcgactatttcggacggc<br>cgtatgaaggcatcgccgtgttcgacggcaaaaagatcactgtaacagggaccctgtggaacggcaacaaaattatcgacgagcgcctgatcaaccccgacg<br>gctccctgctgttccgagtaaccatcaacggagtgaccggctggcggctgtgcgaacgcattctggcgtaagatctttttccctctgccaaaaattatggggacat<br>catgaagccccttgagcatctgacttctggctaataaaggaaatttattttcattgc |
| **Weak CUG (−3C/+4C)** | acacttgcttttgacacaactgtgtttacttgcaatcccccaaaacagccactgcatcatcatcatcatcatggctcgagcggcgtcttcacactcgaagatttcgttg<br>gggactggcgacagacagccggctacaacctggaccaagtccttgaacagggaggtgtgtccagtttgtttcagaatctcggggtgtccgtaactccgatccaa<br>aggattgtcctgagcggtgaaaatgggctgaagatcgacatccatgtcatcatcccgtatgaaggtctgagcggcgaccaaatgggccagatcgaaaaaattttt<br>aaggtggtgtaccctgtggatgatcatcactttaaggtgatcctgcactatggcacactggtaatcgacggggttacgccgaacatgatcgactatttcggacggc<br>cgtatgaaggcatcgccgtgttcgacggcaaaaagatcactgtaacagggaccctgtggaacggcaacaaaattatcgacgagcgcctgatcaaccccgacg<br>gctccctgctgttccgagtaaccatcaacggagtgaccggctggcggctgtgcgaacgcattctggcgtaagatctttttccctctgccaaaaattatggggacat<br>catgaagccccttgagcatctgacttctggctaataaaggaaatttattttcattgc |
| **c-Myc-NLuc (WT)** | aactcgctgtagtaattccagcgagaggcagagggagcgagcgggcggccggctagggtggaagagccgggcgagcagagctgcgctgcgggcgtcctg<br>ggaagggagatccggagcgaataggggggcttcgcctctggcccagccctcccgctgatcccccagccagcggtccgcaacccttgccgcatccacgaaactt<br>tgcccatagcagcgggcgggcactttgcactggaacttacaacacccgagcaaggacgcgactctcccgacgcggggaggctattctgcccatttggggaca<br>cttccccgccgctgccaggacccgcttctctgaaaggctctccttgcagctgcttagacgctggatttttttcgggtagtggaaaaccagcagcctcccgcgacga<br>tgcccctcaacgttagcttcaccaacaggaactatgacctcgactacggctcgagcggcgactataaggaccacgacggagactacaaggatcatgatattgatt<br>acaaggacgacgatgacaagggaggaggaggaagcgtcttcacactcgaagatttcgttggggactggcgacagacagccggctacaacctggaccaagtc<br>cttgaacagggaggtgtgtccagtttgtttcagaatctcggggtgtccgtaactccgatccaaaggattgtcctgagcggtgaaaatgggctgaagatcgacatcc<br>atgtcatcatcccgtatgaaggtctgagcggcgaccaaatgggccagatcgaaaaaatttttaaggtggtgtaccctgtggatgatcatcactttaaggtgatcctg<br>cactatggcacactggtaatcgacggggttacgccgaacatgatcgactatttcggacggccgtatgaaggcatcgccgtgttcgacggcaaaaagatcactgt<br>aacagggaccctgtggaacggcaacaaaattatcgacgagcgcctgatcaaccccgacggctccctgctgttccgagtaaccatcaacggagtgaccggctg<br>gcggctgtgcgaacgcattctggcgtaagatctttttccctctgccaaaaattatggggacatcatgaagccccttgagcatctgacttctggctaataaaggaaatt<br>tattttcattgc |
| **c-Myc-NLuc (Mut 1)** | aactcgctgtagtaattccagcgagaggcagagggagcgagcgggcggccggctagggtggaagagccgggcgagcagagctgcgctgcgggcgtcctg<br>ggaagggagatccggagcgaataggggggcttcgcctctggcccagccctcccgctgatcccccagccagcggtccgcaacccttgccgcatccacgaaactt<br>tgcccatagcagcgggcgggcactttgcactggaacttacaacacccgagcaaggacgcgactctcccgacgcggggaggctattctgcccatttggggaca<br>cttccccgccgctgccaggacccgcttctctgaaaggctctccttgcagctgcttagacgccggatttttttcgggtagtggaaaaccagcagcctcccgcgacg<br>atgcccctcaacgttagcttcaccaacaggaactatgacctcgactacggctcgagcggcgactataaggaccacgacggagactacaaggatcatgatattga<br>ttacaaggacgacgatgacaagggaggaggaggaagcgtcttcacactcgaagatttcgttggggactggcgacagacagccggctacaacctggaccaagt<br>ccttgaacagggaggtgtgtccagtttgtttcagaatctcggggtgtccgtaactccgatccaaaggattgtcctgagcggtgaaaatgggctgaagatcgacatc<br>catgtcatcatcccgtatgaaggtctgagcggcgaccaaatgggccagatcgaaaaaatttttaaggtggtgtaccctgtggatgatcatcactttaaggtgatcct<br>gcactatggcacactggtaatcgacggggttacgccgaacatgatcgactatttcggacggccgtatgaaggcatcgccgtgttcgacggcaaaaagatcactg |

| | |
|---|---|
| | taacagggaccctgtggaacggcaacaaaattatcgacgagcgcctgatcaaccccgacggctccctgctgttccgagtaaccatcaacggagtgaccggctg<br>gcggctgtgcgaacgcattctggcgtaagatctttttccctctgccaaaaattatggggacatcatgaagccccttgagcatctgacttctggctaataaaggaaatt<br>tattttcattgc |
| **c-Myc-NLuc (Mut 2)** | aactcgctgtagtaattccagcgagaggcagagggagcgagcgggcggccggctagggtggaagagccgggcgagcagagctgcgctgcgggcgtcctg<br>ggaagggagatccggagcgaataggggggcttcgcctctggcccagccctcccgctgatcccccagccagcggtccgcaacccttgccgcatccacgaaactt<br>tgcccatagcagcgggcgggcactttgcactggaacttacaacacccgagcaaggacgcgactctcccgacgcggggaggctattctgcccatttggggaca<br>cttccccgccgctgccaggacccgcttctctgaaaggctctccttgcagctgcttagacgctggatttttttcgggtagtggaaaaccagcagcctcccgcgacga<br>gacccctcaacgttagcttcaccaacaggaactatgacctcgactacggctcgagcggcgactataaggaccacgacggagactacaaggatcatgatattgat<br>tacaaggacgacgatgacaagggaggaggaggaagcgtcttcacactcgaagatttcgttggggactggcgacagacagccggctacaacctggaccaagt<br>ccttgaacagggaggtgtgtccagtttgtttcagaatctcggggtgtccgtaactccgatccaaaggattgtcctgagcggtgaaaatgggctgaagatcgacatc<br>catgtcatcatcccgtatgaaggtctgagcggcgaccaaatgggccagatcgaaaaaatttttaaggtggtgtaccctgtggatgatcatcactttaaggtgatcct<br>gcactatggcacactggtaatcgacggggttacgccgaacatgatcgactatttcggacggccgtatgaaggcatcgccgtgttcgacggcaaaaagatcactg<br>taacagggaccctgtggaacggcaacaaaattatcgacgagcgcctgatcaaccccgacggctccctgctgttccgagtaaccatcaacggagtgaccggctg<br>gcggctgtgcgaacgcattctggcgtaagatctttttccctctgccaaaaattatggggacatcatgaagccccttgagcatctgacttctggctaataaaggaaatt<br>tattttcattgc |
| **c-Myc-NLuc (Mut 3)** | aactcgctgtagtaattccagcgagaggcagagggagcgagcgggcggccggctagggtggaagagccgggcgagcagagctgcgctgcgggcgtcctg<br>ggaagggagatccggagcgaataggggggcttcgcctctggcccagccctcccgctgatcccccagccagcggtccgcaacccttgccgcatccacgaaactt<br>tgcccatagcagcgggcgggcactttgcactggaacttacaacacccgagcaaggacgcgactctcccgacgcggggaggctattctgcccatttggggaca<br>cttccccgccgctgccaggacccgcttctctgaaaggctctccttgcagctgcttagacgctggatttttttcgggtagaggaaaaccagcagcctcccgcgacg<br>agacccctcaacgttagcttcaccaacaggaactatgacctcgactacggctcgagcggcgactataaggaccacgacggagactacaaggatcatgatattg<br>attacaaggacgacgatgacaagggaggaggaggaagcgtcttcacactcgaagatttcgttggggactggcgacagacagccggctacaacctggaccaa<br>gtccttgaacagggaggtgtgtccagtttgtttcagaatctcggggtgtccgtaactccgatccaaaggattgtcctgagcggtgaaaatgggctgaagatcgaca<br>tccatgtcatcatcccgtatgaaggtctgagcggcgaccaaatgggccagatcgaaaaaatttttaaggtggtgtaccctgtggatgatcatcactttaaggtgatc<br>ctgcactatggcacactggtaatcgacggggttacgccgaacatgatcgactatttcggacggccgtatgaaggcatcgccgtgttcgacggcaaaaagatcac<br>tgtaacagggaccctgtggaacggcaacaaaattatcgacgagcgcctgatcaaccccgacggctccctgctgttccgagtaaccatcaacggagtgaccggc<br>tggcggctgtgcgaacgcattctggcgtaagatctttttccctctgccaaaaattatggggacatcatgaagccccttgagcatctgacttctggctaataaaggaa<br>atttattttcattgc |
| **c-Myc-NLuc (Mut 4)** | aactcgctgtagtaattccagcgagaggcagagggagcgagcgggcggccggctagggtggaagagccgggcgagcagagctgcgctgcgggcgtcctg<br>ggaagggagatccggagcgaataggggggcttcgcctctggcccagccctcccgctgatcccccagccagcggtccgcaacccttgccgcatccacgaaactt<br>tgcccatagcagcgggcgggcactttgcactggaacttacaacacccgagcaaggacgcgactctcccgacgcggggaggctattctgcccatttggggaca<br>cttccccgccgctgccaggacccgcttctctgaaaggctctccttgcagctgcttagacgctggacttcttccgcgtcgtggagaaccagcagccccccgccac<br>cagacccctcaacgttagcttcaccaacaggaactatgacctcgactacggctcgagcggcgactataaggaccacgacggagactacaaggatcatgatatt<br>gattacaaggacgacgatgacaagggaggaggaggaagcgtcttcacactcgaagatttcgttggggactggcgacagacagccggctacaacctggacca |

| | |
|---|---|
| | agtccttgaacagggaggtgtgtccagtttgtttcagaatctcggggtgtccgtaactccgatccaaaggattgtcctgagcggtgaaaatgggctgaagatcgac<br>atccatgtcatcatcccgtatgaaggtctgagcggcgaccaaatgggccagatcgaaaaaatttttaaggtggtgtaccctgtggatgatcatcactttaaggtgat<br>cctgcactatggcacactggtaatcgacggggttacgccgaacatgatcgactatttcggacggccgtatgaaggcatcgccgtgttcgacggcaaaaagatca<br>ctgtaacagggaccctgtggaacggcaacaaaattatcgacgagcgcctgatcaaccccgacggctccctgctgttccgagtaaccatcaacggagtgaccgg<br>ctggcggctgtgcgaacgcattctggcgtaagatctttttccctctgccaaaaattatggggacatcatgaagccccttgagcatctgacttctggctaataaagga<br>aatttattttcattgc |
| **c-Myc-NLuc (Mut 5)** | aactcgctgtagtaattccagcgagaggcagagggagcgagcgggcggccggctagggtggaagagccgggcgagcagagctgcgctgcgggcgtcctg<br>ggaagggagatccggagcgaataggggggcttcgcctctggcccagccctcccgctgatcccccagccagcggtccgcaacccttgccgcatccacgaaactt<br>tgcccatagcagcgggcgggcactttgcactggaacttacaacacccgagcaaggacgcgactctcccgacgcggggaggctattctgcccatttggggaca<br>cttccccgccgctgccaggacccgcttctctgaaaggctctccttgcagctgcttagacgatggattttttcgggtagtggaaaaccagcagcctcccgcgacga<br>gacccctcaacgttagcttcaccaacaggaactatgacctcgactacggctcgagcggcgactataaggaccacgacggagactacaaggatcatgatattgat<br>tacaaggacgacgatgacaagggaggaggaggaagcgtcttcacactcgaagatttcgttggggactggcgacagacagccggctacaacctggaccaagt<br>ccttgaacagggaggtgtgtccagtttgtttcagaatctcggggtgtccgtaactccgatccaaaggattgtcctgagcggtgaaaatgggctgaagatcgacatc<br>catgtcatcatcccgtatgaaggtctgagcggcgaccaaatgggccagatcgaaaaaatttttaaggtggtgtaccctgtggatgatcatcactttaaggtgatcct<br>gcactatggcacactggtaatcgacggggttacgccgaacatgatcgactatttcggacggccgtatgaaggcatcgccgtgttcgacggcaaaaagatcactg<br>taacagggaccctgtggaacggcaacaaaattatcgacgagcgcctgatcaaccccgacggctccctgctgttccgagtaaccatcaacggagtgaccggctg<br>gcggctgtgcgaacgcattctggcgtaagatctttttccctctgccaaaaattatggggacatcatgaagccccttgagcatctgacttctggctaataaaggaaatt<br>tattttcattgc |
| **c-Myc-NLuc (Mut 6)** | aactcgctgtagtaattccagcgagaggcagagggagcgagcgggcggccggctagggtggaagagccgggcgagcagagctgcgctgcgggcgtcctg<br>ggaagggagatccggagcgaataggggggcttcgcctctggcccagccctcccgctgatcccccagccagcggtccgcaacccttgccgcatccacgaaactt<br>tgcccatagcagcgggcgggcactttgcactggaacttacaacacccgagcaaggacgcgactctcccgacgcggggaggctattctgcccatttggggaca<br>cttccccgccgctgccaggacccgcttctctgaaaggctctccttgcagctgcttagacgccggattttttcgggtagtggaaaaccagcagcctcccgcgacg<br>agacccctcaacgttagcttcaccaacaggaactatgacctcgactacggctcgagcggcgactataaggaccacgacggagactacaaggatcatgatattg<br>attacaaggacgacgatgacaagggaggaggaggaagcgtcttcacactcgaagatttcgttggggactggcgacagacagccggctacaacctggaccaa<br>gtccttgaacagggaggtgtgtccagtttgtttcagaatctcggggtgtccgtaactccgatccaaaggattgtcctgagcggtgaaaatgggctgaagatcgaca<br>tccatgtcatcatcccgtatgaaggtctgagcggcgaccaaatgggccagatcgaaaaaatttttaaggtggtgtaccctgtggatgatcatcactttaaggtgatc<br>ctgcactatggcacactggtaatcgacggggttacgccgaacatgatcgactatttcggacggccgtatgaaggcatcgccgtgttcgacggcaaaaagatcac<br>tgtaacagggaccctgtggaacggcaacaaaattatcgacgagcgcctgatcaaccccgacggctccctgctgttccgagtaaccatcaacggagtgaccggc<br>tggcggctgtgcgaacgcattctggcgtaagatctttttccctctgccaaaaattatggggacatcatgaagccccttgagcatctgacttctggctaataaaggaa<br>atttattttcattgc |
| **c-Myc-NLuc (Mut 7)** | aactcgctgtagtaattccagcgagaggcagagggagcgagcgggcggccggctagggtggaagagccgggcgagcagagctgcgctgcgggcgtcctg<br>ggaagggagatccggagcgaataggggggcttcgcctctggcccagccctcccgctgatcccccagccagcggtccgcaacccttgccgcatccacgaaactt<br>tgcccatagcagcgggcgggcactttgcactggaacttacaacacccgagcaaggacgcgactctcccgacgcggggaggctattctgcccatttggggaca |

| | |
|---|---|
| | cttccccgccgctgccaggacccgcttctctgaaaggctctccttgcagctgcttagacgccggattttttcgggtagaggaaaaccagcagcctcccgcgacg<br>atgcccctcaacgttagcttcaccaacaggaactatgacctcgactacggctcgagcggcgactataaggaccacgacggagactacaaggatcatgatattga<br>ttacaaggacgacgatgacaagggaggaggaggaagcgtcttcacactcgaagatttcgttggggactggcgacagacagccggctacaacctggaccaagt<br>ccttgaacagggaggtgtgtccagtttgtttcagaatctcggggtgtccgtaactccgatccaaaggattgtcctgagcggtgaaaatgggctgaagatcgacatc<br>catgtcatcatcccgtatgaaggtctgagcggcgaccaaatgggccagatcgaaaaaatttttaaggtggtgtaccctgtggatgatcatcactttaaggtgatcct<br>gcactatggcacactggtaatcgacggggttacgccgaacatgatcgactatttcggacggccgtatgaaggcatcgccgtgttcgacggcaaaaagatcactg<br>taacagggaccctgtggaacggcaacaaaattatcgacgagcgcctgatcaaccccgacggctccctgctgttccgagtaaccatcaacggagtgaccggctg<br>gcggctgtgcgaacgcattctggcgtaagatctttttccctctgccaaaaattatggggacatcatgaagccccttgagcatctgacttctggctaataaaggaaatt<br>tattttcattgc |

**Color code for NLuc constructs:** green, *β-globin* 5′ and 3′ untranslated regions (UTRs) (*Oryctolagus cuniculus*; NM_001082260.3); red, start codon (Kozak context); brown, GSSG linker; blue, NanoLuc (NLuc) coding sequence (pNL1.1 [NLuc]; Promega).

**Color code for *c-Myc* constructs:** black, 5′ untranslated region (UTR) of *c-MYC* (*Homo sapiens*; NM_002467.6); red, start codons (common codons); purple, *c-MYC* coding sequence (*Homo sapiens*; NM_002467.6); brown, GSSG linker; green, 3×FLAG tag; blue, NanoLuc (NLuc) coding sequence (pNL1.1 [NLuc]; Promega); gray, *β-globin* 3′ untranslated region (UTR) (*Oryctolagus cuniculus*; NM_001082260.
